# Learning fast and slow: deviations from the matching law can reflect an optimal strategy under uncertainty

**DOI:** 10.1101/141309

**Authors:** Kiyohito Iigaya, Yashar Ahmadian, Leo P. Sugrue, Greg S. Corrado, Yonatan Loewenstein, William T. Newsome, Stefano Fusi

## Abstract

Behavior which deviates from our normative expectations often appears irrational. A classic example concerns the question of how choice should be distributed among multiple alternatives. The so-called matching law predicts that the fraction of choices made to any option should match the fraction of total rewards earned from the option. This choice strategy can maximize reward in a stationary reward schedule. Empirically, however, behavior often deviates from this ideal. While such deviations have often been interpreted as reflecting ‘noisy’, suboptimal, decision-making, here we instead suggest that they reflect a strategy which is adaptive in nonstationary and uncertain environments. We analyze the results of a dynamic foraging task. Animals exhibited significant deviations from matching, and animals turned out to be able to collect more rewards when deviation was larger. We show that this behavior can be understood if one considers that animals had incomplete information about the environments dynamics. In particular, using computational models, we show that in such nonstationary environments, learning on both fast and slow timescales is beneficial. Learning on fast timescales means that an animal can react to sudden changes in the environment, though this inevitably introduces large fluctuations (*variance*) in value estimates. Concurrently, learning on slow timescales reduces the amplitude of these fluctuations at the price of introducing a *bias* that causes systematic deviations. We confirm this prediction in data – monkeys indeed solved the bias-variance tradeoff by combining learning on both fast and slow timescales. Our work suggests that multi-timescale learning could be a biologically plausible mechanism for optimizing decisions under uncertainty.

## 1 Introduction

The matching law constitutes a quantitative description of choice behavior that is often observed in foraging tasks. According to the matching law, subjects distribute their choices across available options in the same proportion as the rewards obtained from those options ([19, 20]). This type of behavior has been observed across a wide range of species including pigeons, rats, monkeys and humans [19, 20, 16, 17, 52, 11, 28, 29, 45, 36, 37]. Although the matching law provides a simple and elegant description of behavior, actual choice often deviated from the strict matching. For example, one common deviation, which is often referred to as *undermatching*, reveals itself as a more random choice allocation, because the subjects systematically choose less-rewarding options more often than the matching law predicts. Such deviations have been sometimes interpreted as a failure of the subjects, which could be caused by poor discrimination between options [3], by noise in the neural mechanisms underlying decision making [50], or by an imbalance in the learning mechanisms [30].

Here we analyzed an experiment in which monkeys were trained to perform a dynamic foraging task [52]. In this task, they had to track the changing values of alternative choices through time. The values (reward rates) changed periodically without any warning. Within each period during which the values were kept constant, the probabilistic strategy that maximizes cumulative reward is to follow the matching law [46, 22]. We show, however, that the animals exhibit a significant deviation from the matching law in the form of undermatching. Intuitively, this deviation from matching behavior should lead to a decreased harvesting performance; but paradoxically we observe that the overall performance increases as the behavior deviates more strongly from the matching law.

This seemingly paradoxical observation is solved if one recognizes that the reward rate of each choice changes unpredictably in our task. Under such uncertainty, the animal has to continuously update the choice values by integrating reward history. In fact, the matching law has been proven to be the stochastic strategy that maximizes the average reward only when the environment is stationary [46, 22]. In a non-stationary situation, such as in our task, the animals should properly weight their past experiences to make a flexible decision, possibly in a way that depends on the degree of uncertainty or volatility of the environment [5, 35, 21]. Indeed, if the environment is stable, it is beneficial to consider a large number of past experiences to better estimate the value of competing alternative choices. If the environment is volatile, then the animal should consider only a relatively small number of recent experiences, as old ones may no longer be informative about the current choice values.

As a consequence there is an optimal way of weighting recent and old experiences. Old experiences, when considered, should introduce a bias away from behavior that optimally follows only the true current choice values (which are, however, unknown to the subject). This bias contains information about the long-run average of true choice values. When different choices are equally rewarded in the long-run, the bias will be toward balanced choices. This in turn means that subjects would choose the options with lower reward rates more often than they should, if they knew the true rates. In our experiment, this bias translates into undermatching; choice behavior appears to be more random and exploratory when compared to matching behavior, which would be optimal for a subject who knew the true reward probabilities. However, the bias is also accompanied by a reduction in the fluctuations in the estimate of current reward probabilities which are inferred from finite stochastic observations. This reduction in the fluctuations amply compensates for deviation from matching behavior. Thus the overall harvesting performance can increase when more old experiences are taken into account. This simple argument shows that the observed deviations from the matching law might actually be a consequence of a more sophisticated adaptive strategy that involves learning over a wide range of timescales.

To test this hypothesis, we estimated the time over which monkeys were integrating the rewards received for each choice. We found that this integration-time, which turned out to be much longer than what has normally been studied, varied slowly across days of experiments, and that it correlated with undermatching. More specifically, larger deviations from matching behavior were observed to correlate with longer integration-times, as we predicted. We also observed that longer integration-times correspond to smaller fluctuations in choice behavior, which reflect a reduced variance in estimate of the choice value. This reduction in the fluctuations is what explains the improved overall performance, and indeed, the harvesting performance increases when the fluctuations decrease.

## 2 Results

### 2.1 The dynamic foraging task

On each trial, the monkey is free to choose between two color targets by making saccadic movements (see Figure 1a). Rewards are assigned to the two colors randomly, at rates that remain constant for a certain number of trials (block size: typically 100–200 trials). Once the reward is assigned to a target, the target is said to be *baited*, and the reward remains available until the target is chosen. This means that the probability of being rewarded from a target increases with the time since the target was last chosen. In a stationary environment under this reward schedule, matching is known to be the probabilistic strategy that maximizes the average chance of obtaining rewards. In that sense matching could be considered as “optimal” (see also [31]). However, in this task the reward rates were periodically modified in a random and unpredictable way.

**Figure 1:**
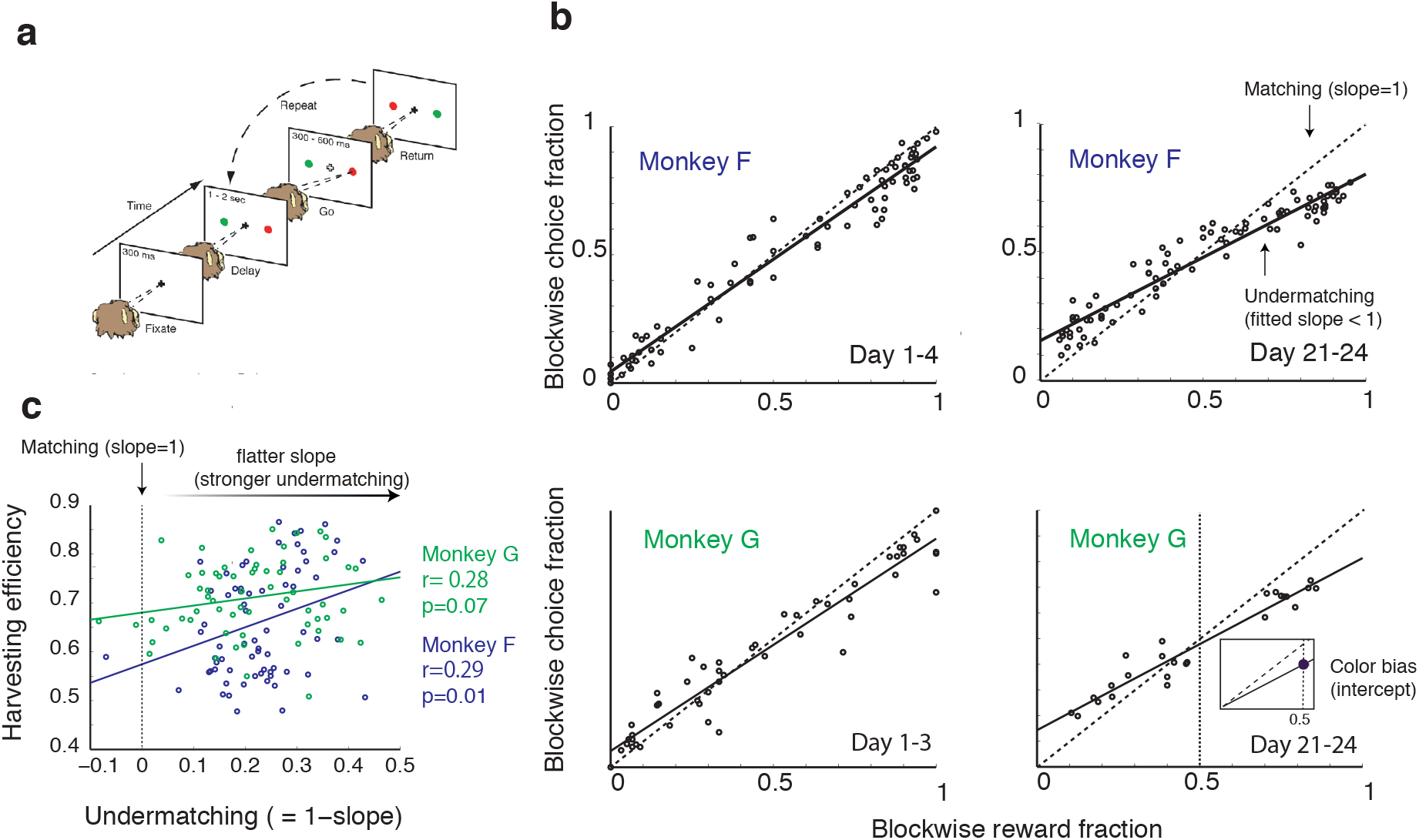
(**a**). Behavioral protocol: the animal had to fixate the central cross, and after a short delay (Delay), it could make a saccadic eye movement toward one of the color targets (Go). If the chosen target was baited, a drop of water was delivered (Return). A delivery of reward resets the target to be empty, until it is baited again, which was stochastically determined with different baiting rates for different targets. The overall maximum baiting rate was set at about 0.35 rewards per trial. The relative baiting rates changed at the end of blocks (about every 100 trials) with no signal. The Ratio of baiting rates in each block was chosen unpredictably from the set (8:1, 6:1, 3:1, 1:1). In this setup if the ratio is fixed, the matching law is known to approximate the optimal stochastic choice behavior. (**b**). Deviation from the matching law: the fraction of choices allocated to one target is plotted as a function of the fraction of rewards that were obtained from the same target for different experimental days (top left Monkey F day 1–4, bottom left: day 21–24, top right Monkey G day 1–3, bottom right: day 21–24). Each data point represents an estimate in a given block of trials, the solid line is a linear fit to the data. The matching law corresponds to a line with a slope equal to 1 (dashed line), while the observed behavior, with a slope < 1, is called undermatching. Undermatching indicates that animals had a tendency to explore choices more (or, put simply, appear to be more random) than what the matching law would predict. For both monkeys the behavior deviates from the matching law, and the degree of undermatching (measured by the slope) changes over time. Note that undermatching is different from color bias, which is indicated by the filled circle in the inner panel (bottom right). The color bias is defined by the intercept of the fitted matching slope and the reward fraction of 0.5. (**c**). Paradoxically, the harvesting efficiency, which indicates how well the monkeys collected rewards, positively correlates with the degree of undermatching: the larger choice behavior deviates from the matching law, the higher the harvesting efficiency. The harvesting efficiency is defined as the number of rewards that monkeys actually obtained divided by the maximum number of rewards that could have been collected. Hence it varies between 0 and 1. The monkeys almost always undermatched, the degree of which shows a wide distribution over sessions.

Nonetheless, the matching law describes fairly accurately the behavior of both monkeys, as already reported in [52]. Now we plotted the fraction of times the monkeys choose one target versus the fraction of times that target was rewarded in Figure 1b. All datapoints are around the diagonal (blue). However, there are clear deviations from the matching law, which become even more evident by comparing a linear fit (red line) of the datapoints to the diagonal. This is a signature of widely observed undermatching, as the choices of the animals tend to be closer to indifferent (choice fraction close to 0.5) than what is predicted by the matching law.

There are now two important observations to be made: the first one is that deviation from the matching law seems to vary over time (see for example deviations estimated over two time intervals in Figure 1b that are clearly different). One way to express more quantitatively this deviation is to compute the slope S of the best linear fit and compare it to the unitary slope of the diagonal. More specifically, we will express the degree of undermatching as 1 − *S*. We observed by analyzing the data that this quantity varies significantly over time, ranging mostly from 0.1 to 0.4.

The second observation is that the slope changes are accompanied by changes in the overall performance expressed as harvesting efficiency (the number of rewards that subjects actually obtained divided by the maximum number of rewards that could have been collected). Paradoxically, the harvesting efficiency increases as the behavior deviates more prominently from the matching law (see Figure 1c). This observation seems to be incompatible with the statement that matching behavior is optimal in this task. However, as we will explain in the next section using computational models, it is actually what we should have expected in a non-stationary environment in which the reward probabilities change over time.

Note that there is another well-observed deviation from the matching law [3], which we refer to as *color bias*. This is a bias toward one of the colored targets, and we express it more quantitatively as the value of the linear fit at 0.5 reward fraction (see the inner panel of Figure 1b and Method section). It is important to distinguish the two biases: undermatching and color bias, as these are independent measures. In what follows we will focus on undermatching.

### 2.2 The bias-variance tradeoff and the expected changes in matching behavior

Consider the dynamic foraging task that we analyzed. One way of modeling the decision process is to integrate rewards for each choice over a certain number of trials. As we discussed in the Introduction, in a non-stationary situation in which the reward probabilities change from time to time, subjects should properly weight their past experiences, possibly in a way that depends on the volatility of the environment, which in our case is related to how often the reward probabilities change. More specifically, in our case the reward probabilities are constant in each block of 100–200 trials. When the reward probabilities do not change or change rarely, subjects should consider a large number of past experiences to improve the estimate of the reward probabilities. However, if reward probabilities change often, then the subjects should consider only the recent experiences that reflect the current choice values. One simple way of changing the relative weights of recent and remote experiences is to consider multiple exponential integrators [11], each one integrating the reward streams for a specific target on a different timescale.

The mechanism is described schematically in Figure 2a. Consider the case of two exponential integrators characterized by two time constants *τ_Fast_*, *τ_Slow_*. There are two integrators (slow and fast) per choice, and each integrator is represented in the figure by a box. The two top ones integrate the reward stream for the green target, whereas the two bottom ones integrate the reward stream for the other, red, target. The integrator outputs are basically estimates of the value of a particular choice based on a certain number of recent experiences, which is determined by the time constant *τ_Fast_* or *τ_Slow_*. We define the local income [52] of each target to be a weighted average of the output of the fast and slow integrators for that target.

**Figure 2:**
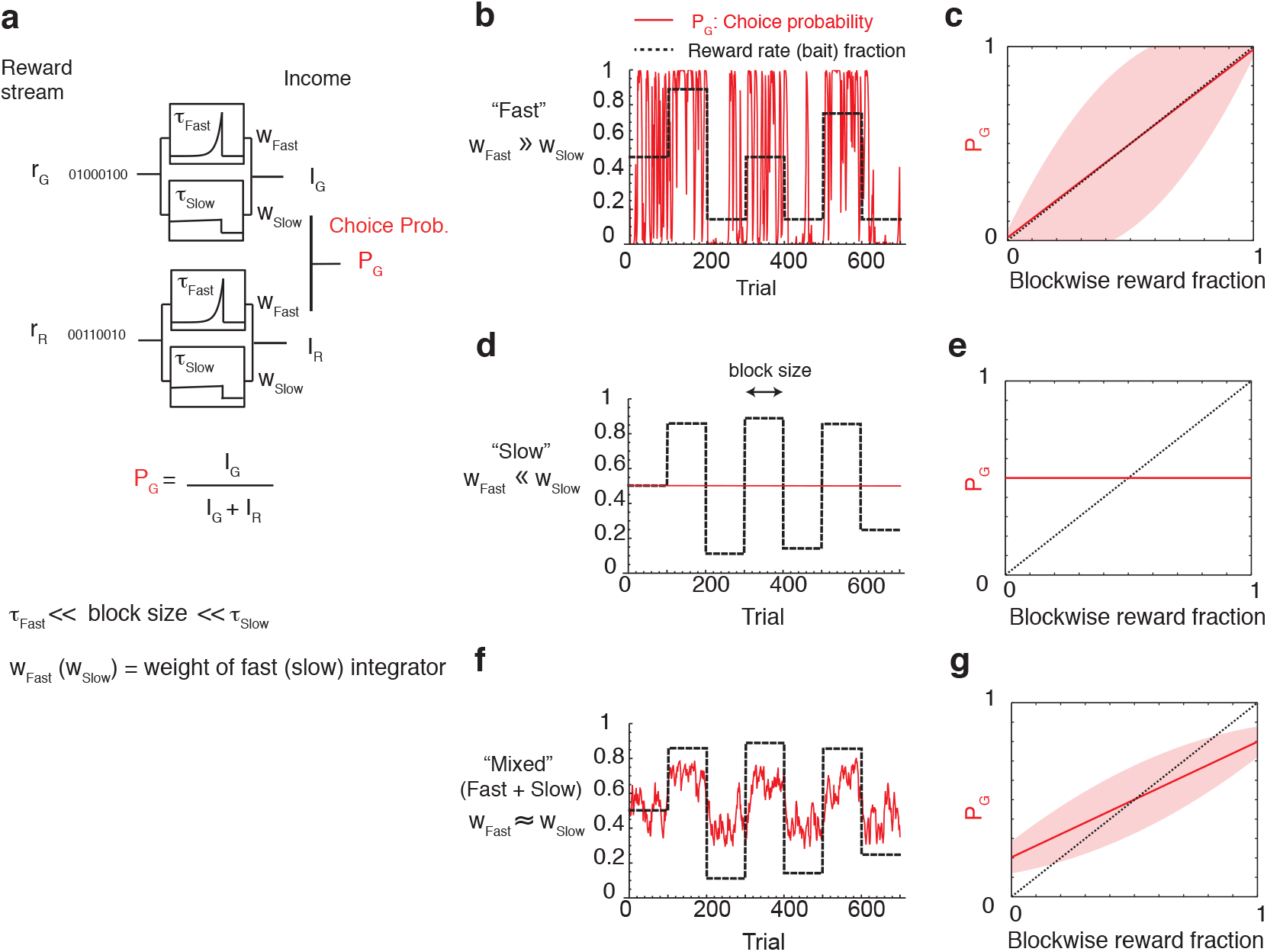
Explaining how deviation from the matching law can improve performance. (**a**) Scheme of a simple decision-making model that is known from previous works to reproduce the behavior observed in the experiment. The model integrates reward history over two timescales (*τ_Fast_*, *τ_Slow_*) to estimate the expected income for each choice (red or green). These incomes are then combined together to generate stochastic decisions, where *P_G_* (*P_R_* = 1 − *P_G_*) is the probability of choosing the green (red) target. While the previous models focused on the integration timescales that are shorter than the block size [52, 11], here we assume that the long timescale τ_Slow_ is much longer than the block size, while the short timescale *τ_Fast_* is still shorter than the block size. The model weights differently the incomes estimated over the two timescales. More specifically *w_Fast_* is the relative weight of the fast integrator (*τ_Fast_*) and *w_Slow_* the weight of the slow one (*w_Fast_* + *w_Slow_* = 1). (**b,c**) Fast integration (*w_Fast_* ≫ *w_Slow_*). If the weight of the fast integrator is much larger than the slow one, the model relies only on the recent reward history estimated over an interval that is approximately *τ_Fast_* to make a decision on the choice. As a consequence, the estimated incomes largely and rapidly fluctuate. This noisy estimation leads to large fluctuations in the choice probability *P_G_* (red). Despite the fluctuations, the mean of such choice probability follows the matching law (indicated by the solid red line in (**c**)). However, the fluctuations of *P_G_* are rather large, as indicated by the broad shaded area, which denotes the standard deviation of *P_G_*. (*d,e*) Slow integration (*w_Fast_* ≪ *w_Slow_*). If the weight of the slow integrator is much larger than the fast one, the model now integrates rewards only on the long timescale *τ_Slow_*. This eliminates entirely the fluctuations in the choice probability; however, the probability of choosing any of the two target is constant at 0.5, because the estimated incomes are balanced on average when multiple blocks of trials are considered. As such, the choice probability becomes independent from the recent reward history, causing a strong (exploratory) deviation from the matching law (**e**). Note the the actual choice probability is determined by the overall reward-color bias in the task (0.5, if no bias). (**f,g**) Mixed integration: (*w_Fast_* ≃ *w_Slow_*). If the two integrators are almost equally weighted, both deviation from the matching law (undermatching) and the amplitude of the fluctuations are intermediate. This matches with observed data, and manifests a computational tradeoff between bias (long integrator; undermatching) and variance (short integrator; fluctuations). Parameters were set to be *τ_Fast_* = 5 trials, *τ_Slow_* = 10,000 trials, *w_Slow_* = 0.3 for (**f,g**). Note that our results do not rely on the specific choice of *τ_Fast_* and *τ_Slow_*.

The decision is then assumed to be the result of the comparison between the local incomes for the two targets. Following [52], the choices are stochastic with a probability of choosing the green target that is given by:

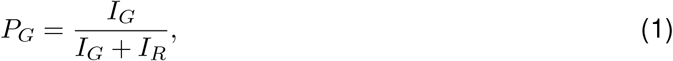

where *I_G_/_R_* is the local income for Green/Red target. While this is not the only way of modeling decisions that depend on past experiences, it is simple to analyze and it reproduces the animals’ behavior with a reasonable approximation [52, 11]. Note, though, that in the previous analysis only short timescales (*τ_Fast_*) have been considered for this model.

The statistics of the decisions generated by the model clearly depend on the timescales *τ_Fast_* and *τ_Slow_*. Consider the case in which *τ_Fast_* is short (shorter than the typical block lengths if expressed in number of trials), and *τ_Slow_* is very long (longer than the typical block lengths), so that the second integrator with *τ_Slow_* integrates the reward streams over multiple blocks of trials. If the weight *w_Fast_* of the first exponential integrator is much larger than *w_Slow_* (Figure 2b-c), then the average choice fraction rapidly tracks the recent average reward fraction. Fast learning is especially advantageous when adapting to a rapid change in the reward contingency at the block transitions (Figure 2b). However, the disadvantage is that the estimated reward fraction can fluctuate wildly, as it follows local noise. This is evident in Figure 2b and it is shown for various reward fractions in Figure 2c, where we plotted the choice fraction vs the reward fraction. In this plot the average (solid line), is very close to the diagonal, indicating that the model has a behavior that follows the block-wise matching law, but the fluctuations are very large (shaded area). The case dominated by *τ_Slow_*, which is the other extreme situation, is illustrated in Figure 2d-e. In this case the integrators estimate the value of each choice over multiple blocks and, as a consequence, the local incomes are constant and equal. This is because the long-run incomes are approximately balanced on average by our experimental design. As a result, the choice shows an extreme undermatching with negligible fluctuations (Figure 2e).

Intermediate situations can be constructed by changing the relative weights of the two integrators. As expected, this model has an intermediate behavior, shown in Figure 2f,g for a particular choice of *w_Fast_*, *w_Slow_*. Increasing the weight of the slow integrator *w_Slow_* would increase deviation from the matching law, which is caused by a bias in the choice value estimates, but it would also decrease the fluctuations. This indicates that the model trades the choice bias (undermatching) for a reduction in the fluctuations, or the variance, of reward estimates by changing the relative contributions of fast and slow integrators.

It is natural to ask whether there is an optimal value of the weight *w_Slow_*. To address this question, we analyzed our model analytically in a more general situation in which the degree of volatility (the block size) is a free parameter (see Supplementary Material section S1 for the calculations). Figures 3a-c show that there is a *w_Slow_* that maximizes the accuracy of the choice value estimate (i.e. it minimizes the mean squared error) and its value does indeed depend on the volatility of the environment (i.e. the number of trials per block). The squared error of the model’s choice value estimate can be expressed as the sum of two terms: the squared bias and the variance. In Figure 3a we show the squared bias and the variance as a function of *w_Slow_*. The variance decreases and the bias increases as *w_Slow_* increases. The mean squared errors of the estimate of the reward probability vs *w_Slow_* is plotted in Figure 2b for two sizes of the blocks of trials in which the reward fraction was constant. Not surprisingly, for more stable environments (dashed line), the optimal weight of long timescales is larger. Consequently, the optimal slope of matching behavior changes according to the volatility (Figure 3c).

**Figure 3:**
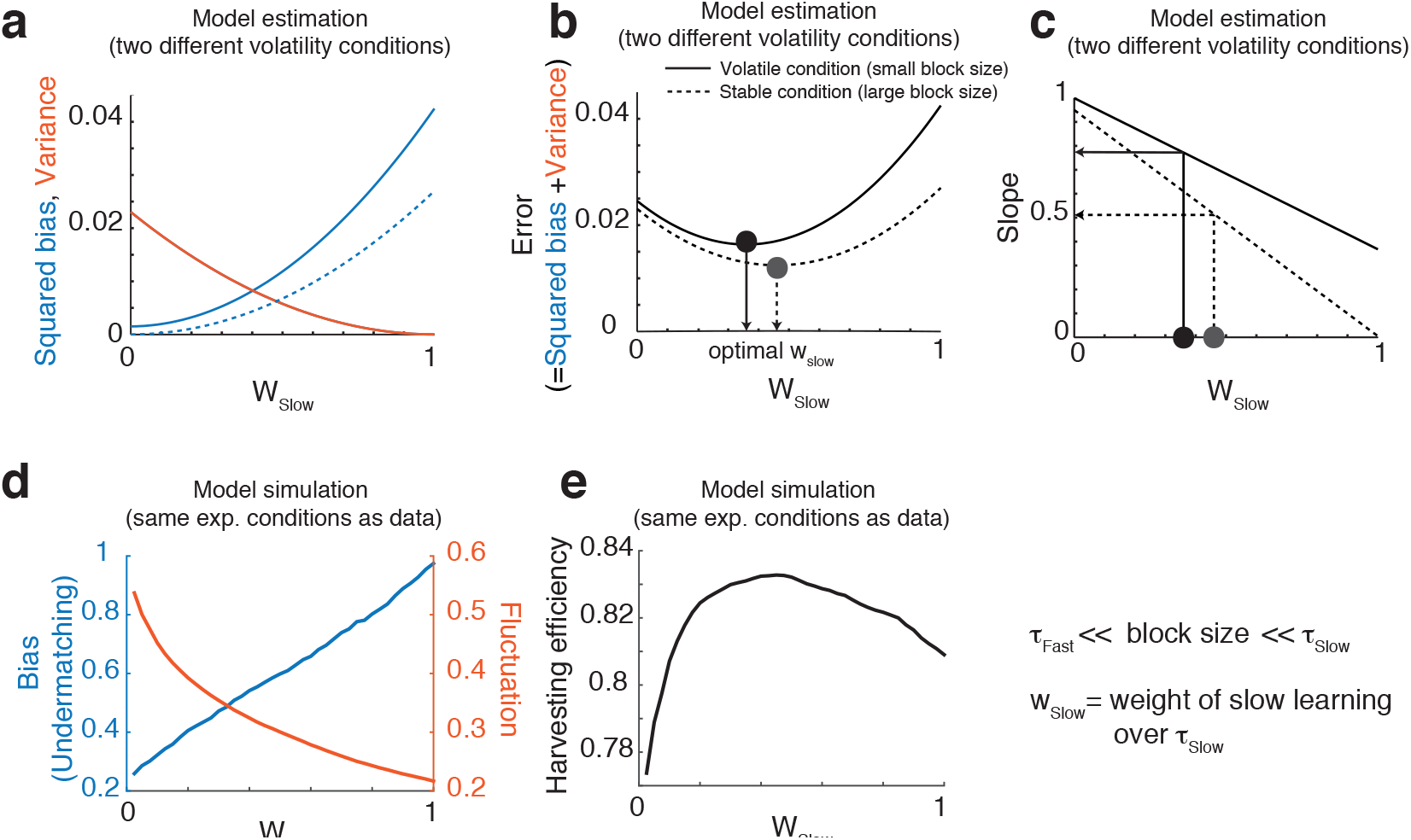
The bias-variance trade-off and the optimal choice behavior under uncertainty, (**a,b,c**) Analytical model results for two different volatility conditions. All the solid (dashed) lines refer to the results for a less (more) volatile task with a block size of 100 (10,000 trials). (**a**) Squared bias (blue) and variance (orange) in the reward rate estimates show a tradeoff as a function of the relative weight of the slow integrator (*w_Slow_*). (**b**). The squared error, which is simply the sum of the squared bias (blue in (**a**)) and the variance (orange in (**a**)), is plotted against *w_Slow_* for different volatility conditions. Intuitively more volatile environments (solid curve) require faster integrators, and hence a smaller relative weight of the slow integrator. Indeed, the optimal relative weight *w_Slow_* for the volatile environment (solid vertical line) is smaller than that for the stable environment (dotted vertical line) (**c**) The deviation from the matching law (here simply shown as the slope of block-wise choice fraction vs reward fraction) also covaries with the relative weight of the slow integrator, and also with the volatility condition. (**d,e**). Model simulation results on the same reward schedule as in the matching experiment. (**d**) Our model simulation also predicts the clear tradeoff between the bias (undermatching) and the fluctuations of choice probability, as a function of the relative weight of slower learning *w_Slow_*. (**e**) Simulation results of harvesting efficiency as a function of the relative weight of the slow integrator *w_Slow_*. As a result of the bias-variance trade-off, the curve takes an inverted U-shape, with a maximum at the optimal relative weight, which is determined by the volatility of the experiment. For panel (**d**), we computed the fluctuation of Monkey’s choice as follows. First, monkey’s choice time series was smoothed via two half-gaussian kernels with standard deviations of σ = 8 trials and *σ* = 50 trials, with a span of 200 trials. This gave us two time series: a fast one with *σ* = 8 and a slow one with *σ* = 50, which we used to compute the bias and variance. We defined the fluctuation as the standard deviation of the fast one over the slow one. The bias was defined by 1– the slope of block-wise matching behavior. For panels (**d,e**) the model with different *w_Slow_* was simulated on Monkey F’s reward schedule. We set *τ_Fast_* = 2, *τ_Slow_* = 1000 trials. Note that our results do not rely on the specific choice of *τ_Fast_* and *τ_Slow_* (please also see Figure S1).

So far we have considered the error on the estimation of the reward probability as a performance measure. In order to evaluate the actual behavioral consequences (e.g. the harvesting efficiency) which will be compared to the experimental data, we simulated our model of Figure 2 for different values of *w_Slow_*, using the same conditions as in the experiments on Monkey F. In Figure 3d we show that undermatching (the bias) is indeed traded off against the amplitude of the fluctuations of the choice probability estimates. A bias of 0 would indicate that the behavior follows the block-wise matching law. As *w_Slow_* increases, undermatching (1– the slope of matching behavior) becomes more prominent, causing the behavior to deviate from strict matching. However, also the fluctuations decrease, as indicated by the red line. Thus as seen in Figure 3e, the overall performance, measured as harvesting efficiency, has a maximum at an intermediate value of *w_Slow_*.

In sum, our computational model analysis suggests that the observed changes in matching behavior could be accounted for by changes in the relative contribution of a (extremely) slow reward integrator to decision making. Moreover, these changes might reflect the bias-variance tradoff in value estimation. In the ideal case, the weight of the fast and slow integrators, *w_Fast_* and *w_Slow_*, should be tuned to the volatility of the environment.

### 2.3 Undermatching observed in the experiments reflects reward history integration over long timescales

Our simple model analysis predicts that deviations from matching behavior observed in the experiments should be related to changes in the relative contribution of the slow integrator. To test this prediction, one could fit the multi-timescale learning model illustrated in Figure 2a to the data, determine how the relative weights of the integrators change over time, and show that these changes correlate with undermatching. We performed this analysis by using the model of Figure 2 with three time constants instead of two. We included two time constants that are shorter than the typical block size, in addition to the one that is significantly longer than the block size. The inclusion of two short time constants, instead of one, was motivated by previous studies on the same experiments [11], which have shown that integrators with two (short) time constants (~ 2 and 20 trials) can describe the animals’ transient behavior better than the integrator with one (short) time constant. Following this finding [11], we set the two short time constants to 2 and 20 trials, and the third one to 1000 trials, so that it spans multiple blocks of trials. Our additional model-agnostic autocorrelation analysis, reported in Figure S2, also provides independent support for such a long time constant (Figure S2b), as well as two short time constants (Figure S2a).

We were surprised to observe unusually smooth changes of the relative weights (*w_Fast_−1*, *w_Fast−2_*, *w_Slow_*) of the three integrators over sessions (Figure 4b), as we did not impose any smoothness constraints to our fitting process (i.e. we fitted each session independently). This suggests that the optimization of relative weights – a type of meta-learning – takes place very slowly, and continuously, over sessions.

**Figure 4:**
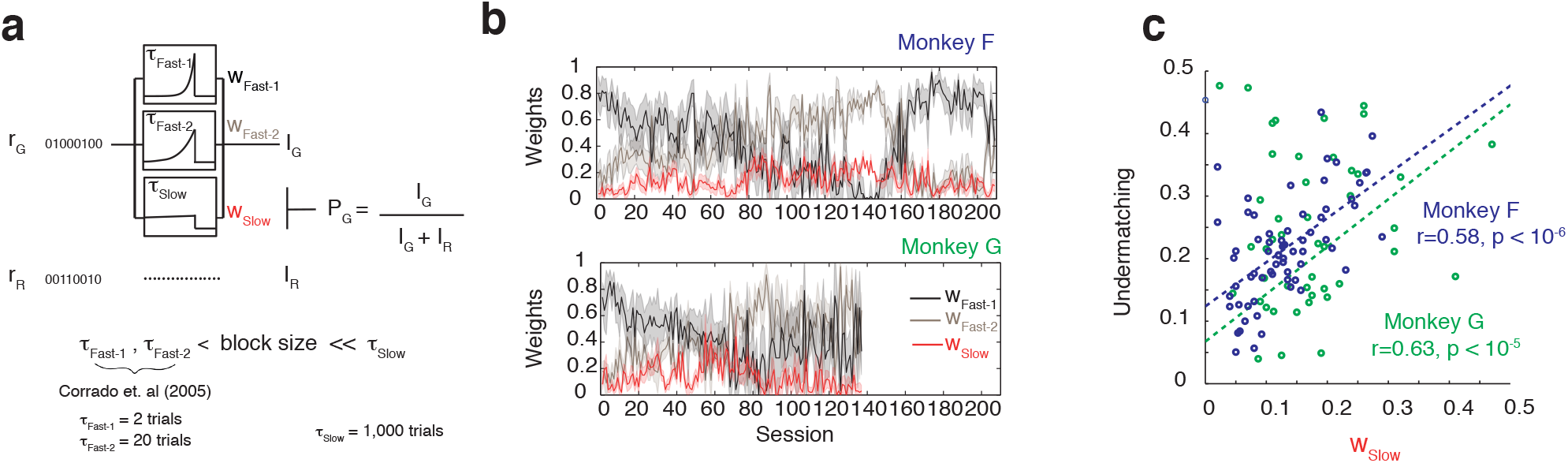
Fitting the multi-timescale model of Figure 2 to the data. (**a**). A model with three timescales (*τ_Fast−1_* = 2 trials, *τ_Fast−2_* =20 trials, and *τ_Slow_* = 1000 trials) is fitted to the data by tuning the weights *w_Fast−1_* (black), *w_Fast−2_* (brown), and *w_Slow_* (red) of the different integrators for each session independently (*w_Fast−1_* + *w_Fast−2_* + *w_Slow_* = 1). (**b**) The weights of different timescales change over consecutive experimental sessions. The short timescales are dominant in early sessions, but the longer timescale becomes progressively more influential. The opposite trend is observed around session 160 of monkey F, probably due to the shortening of experimental sessions and longer inter-experimental-intervals (see Figure S5 for more details). (**c**) Deviation from the matching law is correlated with the weight of the reward integration on a long timescale. Undermatching, computed over the last 50 trials of each block to ignore transients, is plotted against the fitted value of *w_Slow_*, the weight of the longest reward integration timescale. Both monkeys show significant correlations between undermatching (1–slope) and *w_Slow_*. We found, however, this model-based analysis inconclusive, as the correlation is expected from the model structure (please see the text).

Further, as we predicted, changes in the weight of the slow integrator are correlated with changes in undermatching (Figure 4c). However, due to a potential confound, this correlation by itself does not provide conclusive evidence for the hypothesis that undermatching reflects reward history integration over long (but finite) timescales. Indeed, when the time constant of the slow integrator is sufficiently long (compared to experimental block length), a higher *w_Slow_* primarily amplifies the overall degree of undermatching over the entire experimental session, while only negligibly affecting slow adaptive changes in the choice behavior *during* a session.

To understand why, consider the multi-timescale integrator model with two time constants, one fast and one much slower, as in Figure 2, in a volatile environment. The fast integrator will generate estimates of the current values of the two alternatives, which we call 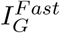 and 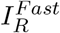 respectively. The slow integrator will generate two additional estimates: 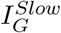 and 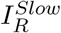, which are almost constant on the timescale of the fast integrator and they act as a bias. Furthermore, because the long-run incomes of the two targets were balanced on average, for *τ_Slow_* much larger than experimental block length, the amount of rewards that animals would receive over this long timescale would be almost the same for the two choices: 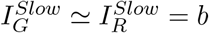, with *b* showing negligible changes over each experimental session. Since the model’s decision policy is described by Equation 1, the probability of choosing *G* will be:

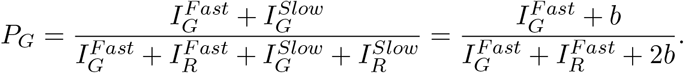

Assuming that the relative weight of the slow integrator is much smaller than the one of the fast integrator (*I_G_* + *I_R_* ≫ 2*b*, or *w_Fast_* ≫ *w_Slow_*), *P_G_* can be expanded as:

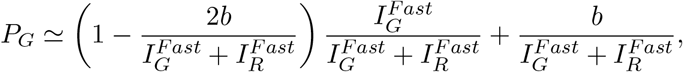

or equivalently

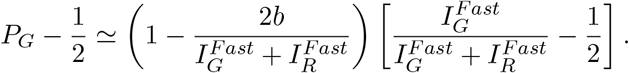

The factor in parenthesis represents a tilt of the curve representing the matching law. In other words, the degree of undermatching is proportional to 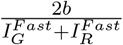, or indeed to *b* itself. As *b* is proportional to the weight of the slow integrator *w_Slow_*, we expect a strong correlation between the fitted weight *w_Slow_* and the average degree of undermatching observed in an experimental session. On the other hand, because b is approximately constant, a higher *w_Slow_* does not necessarily reflect slow adaptive temporal changes during the session. Indeed, even if undermatching had nothing to do with slow reward integration (*e.g.* if it were due to noisy decision making), we can expect that the fit of our model would nevertheless result in higher *w_Slow_* in sessions with stronger undermatching.

Therefore, to address our hypothesis, we need a more direct measure of slow integration times, which does not rely on our model. For this reason, we decided to estimate the lagged correlation between the reward color bias and the choice color bias over multiple experimental sessions. The longest lag for which this correlation is significantly larger than zero provides us with a model-independent (or model-agnostic) estimate of the longest measurable timescale over which the reward bias is estimated. Indeed, a significant correlation would indicate that the choice of the monkey is still affected by the reward bias estimated on a number of trials ago, where the number of trials is determined by the longest lag. Furthermore, this measurement is independent from undermatching, as color bias and undermatching are defined independently.

To assess the validity of this model-independent measure, the correlation is first estimated for the simulated data that are generated by our multi-timescale learning model (the model presented in Figure 2a). Formally, we first estimate reward color bias and choice color bias for each session of simulated experiments. Then we take lagged correlations between these two bias vectors (Figure 5a). Naturally, such correlations decay as the session lag increases. Then we define the longest time lag for which the correlation is significant, which we will call the longest measurable integration timescale (LMIT, see Methods for a more formal definition). For this, we simply fit the correlations linearly, and take the intercept of the linear fit and the 0 correlation. We then transform this session lag measure to trials by multiplying it with the mean session size so that the LMIT is expressed in trials.

**Figure 5:**
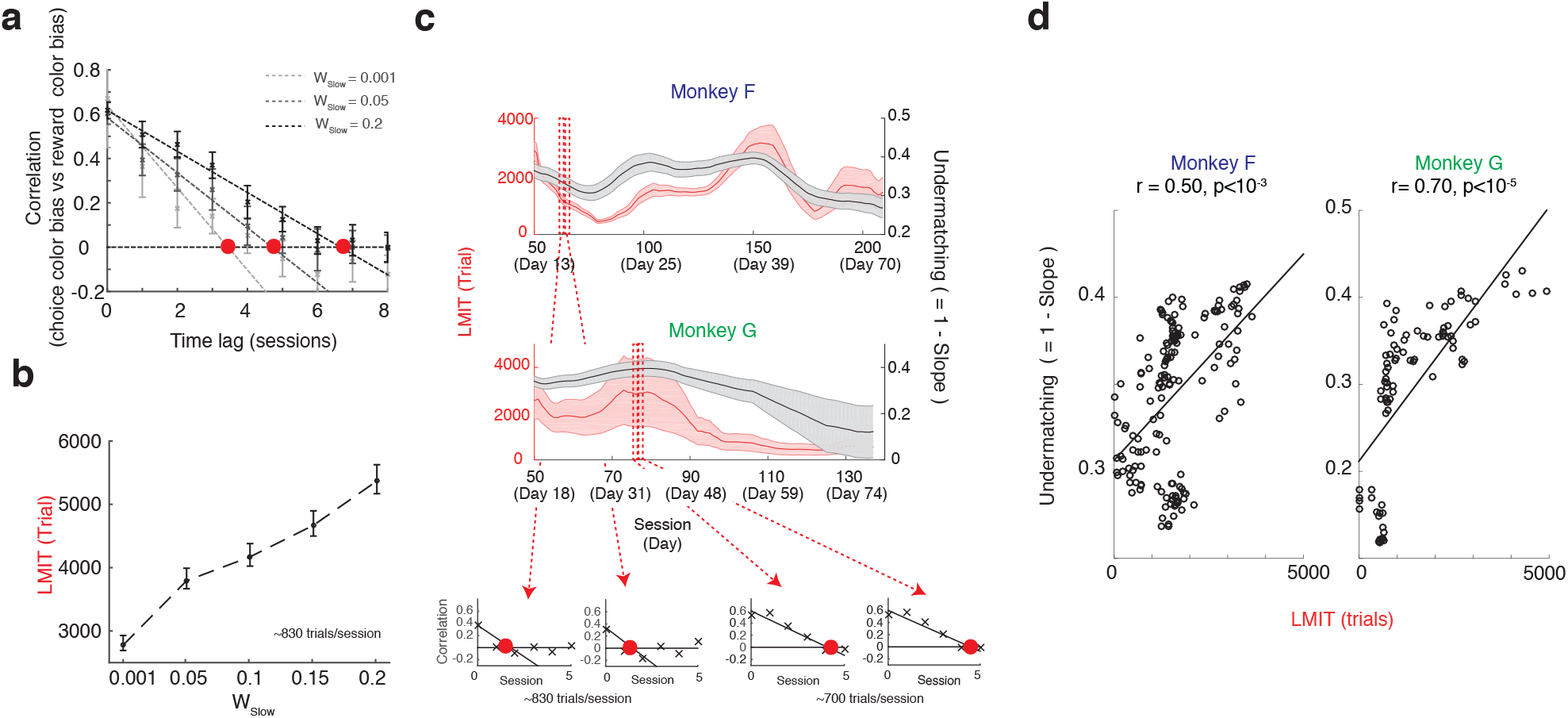
Estimating the longest measurable integration timescale (LMIT) in the model and in the experimental data, and showing how it correlates with undermatching. (**a**). Estimating LMIT in the multi-timescale learning model. Lagged correlation between choice color bias and reward color bias are plotted for different relative weights of the slow integrator *w_Slow_*. Note that these color biases are different from undermatching. The correlation decays as the time-lag between the choice bias and the reward bias increases. The correlation is fitted by a weighted linear regression (dotted lines), and the LMIT is defined by the point at which the fitted line crosses the zero correlation (red filled circles). The correlations are the mean of over 50 simulations and the error bars indicate the standard deviations. The model was simulated over the same reward schedule as Monkey F. The timescales are assumed to be *τ_Fast_* = 5 trials and *τ_Slow_* = 1000 trials, though the specific choice of timescales is not essential for the results. (**b**). The estimated LMIT shows a clear positive correlation with the relative weight of the long timescale *w_Slow_*. The LMIT is expressed in trials, by converting from sessions using the mean session size. This strong monotonic relationship between *w_Slow_* and the LMIT suggests that we can use the LMIT estimate as a proxy for the *w_Slow_* estimate, when analyzing experimental data (see Fig.4 for a direct *w_Slow_* estimation). (**c**). Experimental data: both LMIT (red) and deviation from the matching law (black) vary over time for both monkeys. In the small insets we show four samples of the lagged correlation between choice and reward color bias (the same plot as the one in panel (**a**), but now for the real data). The degree of undermatching seems to be correlated with the LMIT, as predicted by the theory. This impression is confirmed by the analysis in panel (**d**), where we plotted the degree of undermatching as a function of the LMIT. The correlations between the two quantities are highly significant for both monkeys (conservative piece-wise permutation test: *p* < 10^−3^ for monkey F and *p* < 10^−5^ for monkey G). See the main text for the details of the permutation test. See also Figure S3.

The results for our model with different relative weights *w_Slow_* of the longest timescale (*τ_Slow_* = 1000 trials) are plotted as a function of the time lag in Figure 5a. The LMIT is plotted in Figure 5b as a function of the relative weight *w_Slow_*. The LMIT is rather sensitive to changes of *w_Slow_* and hence it is likely that its modifications in time are observable in the behavioral data.

We find that the LMIT indeed covaries with deviation from the matching law. As shown in Figure 5cd for both monkeys, not only the LMIT changes over time (Figure 5c), but it is also correlated with the degree of undermatching (Figure 5d). This confirms our model’s prediction that undermatching (bias towards a 1:1 ratio of choices) reflects very slow reward integrations over hundreds of, or thousands of, trials.

It is important to stress that our predictions did not rely on the details of our computational model (illustrated in the previous section), such as the exact time constants for reward integrations. We conducted a model-agnostic analysis, since we found that fits of our model to data can intrinsically lead to correlation between undermatching and the estimated weight of slow learning. An additional advantage of this model-agnostic approach is that we can test our predictions without determining the parameters of our model, and, even better, without assuming any computational model.

### 2.4 Deviations from the matching law are accompanied by a change of the harvesting performance

Our multi-timescale learning model (Figure 2a) also predicts that deviation from the matching law should also be accompanied by changes in the harvesting efficiency, as a result of the bias-variance tradeoff (see Figure 3e). As the weight of the integrator with the longest timescale increases, the harvesting efficiency should first increase and reach a maximum and then decrease. Hence if w_Slow_, or its proxy – the LMIT, change over time, then it should be possible to observe also changes in the harvesting efficiency. This is indeed the case, as shown in Figure 6a,b.

**Figure 6:**
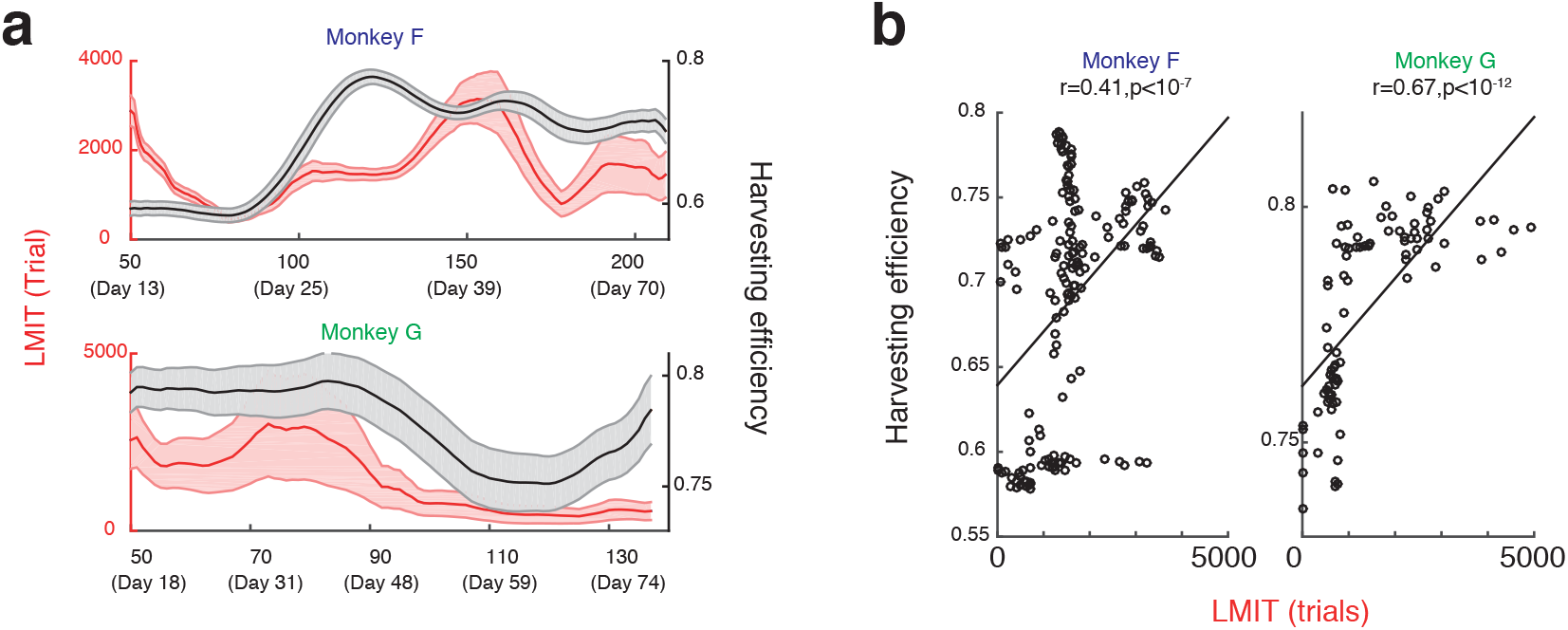
LMIT is also correlated with harvesting efficiency in data, as predicted by our model. (**a**) Changes in LMIT (same as in Figure 5) and harvesting efficiency. The harvesting efficiency was computed for each session, then taken the mean over each reference window to match the estimated LMIT. (**b**) The LMIT and harvesting efficiency show a significant positive correlation for both monkeys, as predicted by our model (The conservative piece-wise permutation test: *p* < 10^−7^ for monkey F and *p* < 10^−12^ for monkey G).

Our explanation for the change in the harvesting efficiency is related to the bias-variance tradeoff. Longer integration-times can reduce the variance but increase the bias. This is what we observed also in the data (see Figure 7a,b). As expected, the bias is negatively correlated with the variance (Figure 7e). Analogously, the harvesting efficiency (Figure 7c,d) is correlated both with the variance (negatively) and the bias (positively) as shown in Figure 7f,g respectively. This is further evidence in support of our model and it explains the seemingly paradoxical effect that monkeys perform better when deviation from the matching law becomes more prominent.

**Figure 7:**
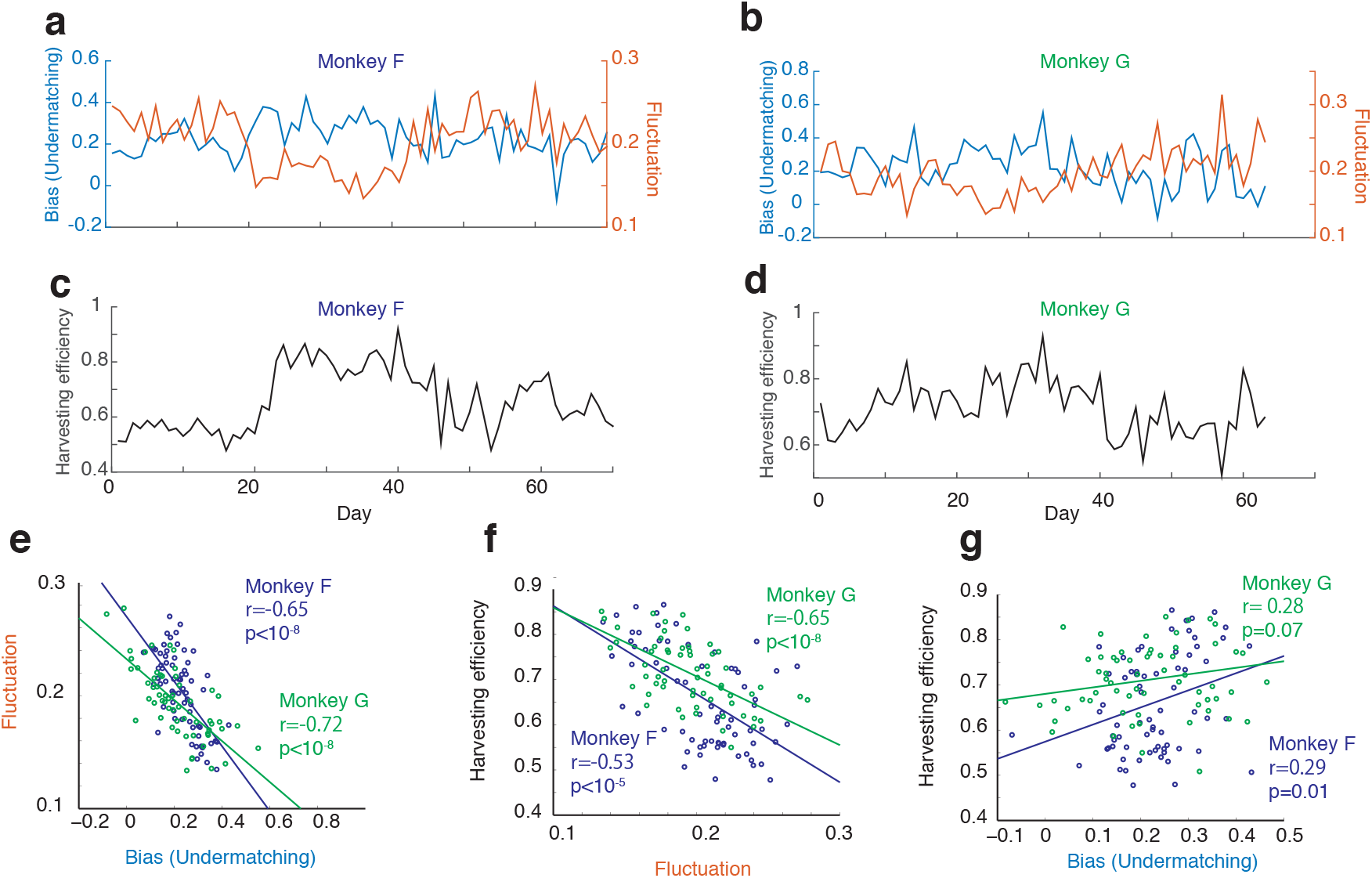
Monkeys show the predicted tradeoff. (**a,b,c,d**) Changes in bias (undermatching), fluctuation and harvesting efficiency over experimental days. (**e,f,g**) Monkeys show the bias-variance tradeoff. (**e**) The bias (1– the slope of matching behavior) and the fluctuation (the square root of variance of monkeys choice probability) are significantly correlated negatively [permutation test: *p* < 10^−8^ for Monkey F and *p* < 10^−8^ for Monkey G]. This means that the monkeys trade off the bias (undermatching) against the variance of estimation, as predicted by our model. (**f**) By reducing fluctuations of estimation, monkeys improved their harvesting performance. The harvesting efficiency is significantly negatively correlated with choice fluctuation [permutation test: *p* < 10^−8^ for Monkey F and *p* < 10^−5^ for Monkey G]. (**g**) Although increasing bias itself should be harmful, the benefit of accompanied reductions in the choice fluctuations lead to a mild increase in the harvesting efficiency [permutation test: *p* < 0.07 for Monkey F and *p* < 0.01 for Monkey G].

It is important to notice that in our model the main reason to modify the relative weights of the integrators is to adapt to the volatility of the environment. In the case of the macaque experiment, there should be no reason for modifying the weights once the optimal value is found. This is because the volatility, determined by the block size, was kept relatively constant. Nevertheless, we found that the relative weights, and analogously the LMIT, changed over the entire course of the experiment (spanned over many days). Explaining these dynamics is beyond the scope of our current model, and it is probably due to factors that are not under control in the experiment. Most likely the weights of the integrators change because of what happens not only during the experiments but also in the intervals between two consecutive experimental sessions. A partial evidence for this explanation is that the LMIT changes in a way that can be at least partially related to the interval between recent experiments and the duration of consecutive experimental sessions (see Figure 8). Specifically, we found that the LMIT decreased when the mean recent intervals between experiments (mean recent break length) were long. On the contrary, the LMIT increased when recent experimental sessions were longer and contained more trials (mean recent experimental session length). This is also the case for the relative weight of slow learning of our model that is fitted to data (Figure S5).

**Figure 8:**
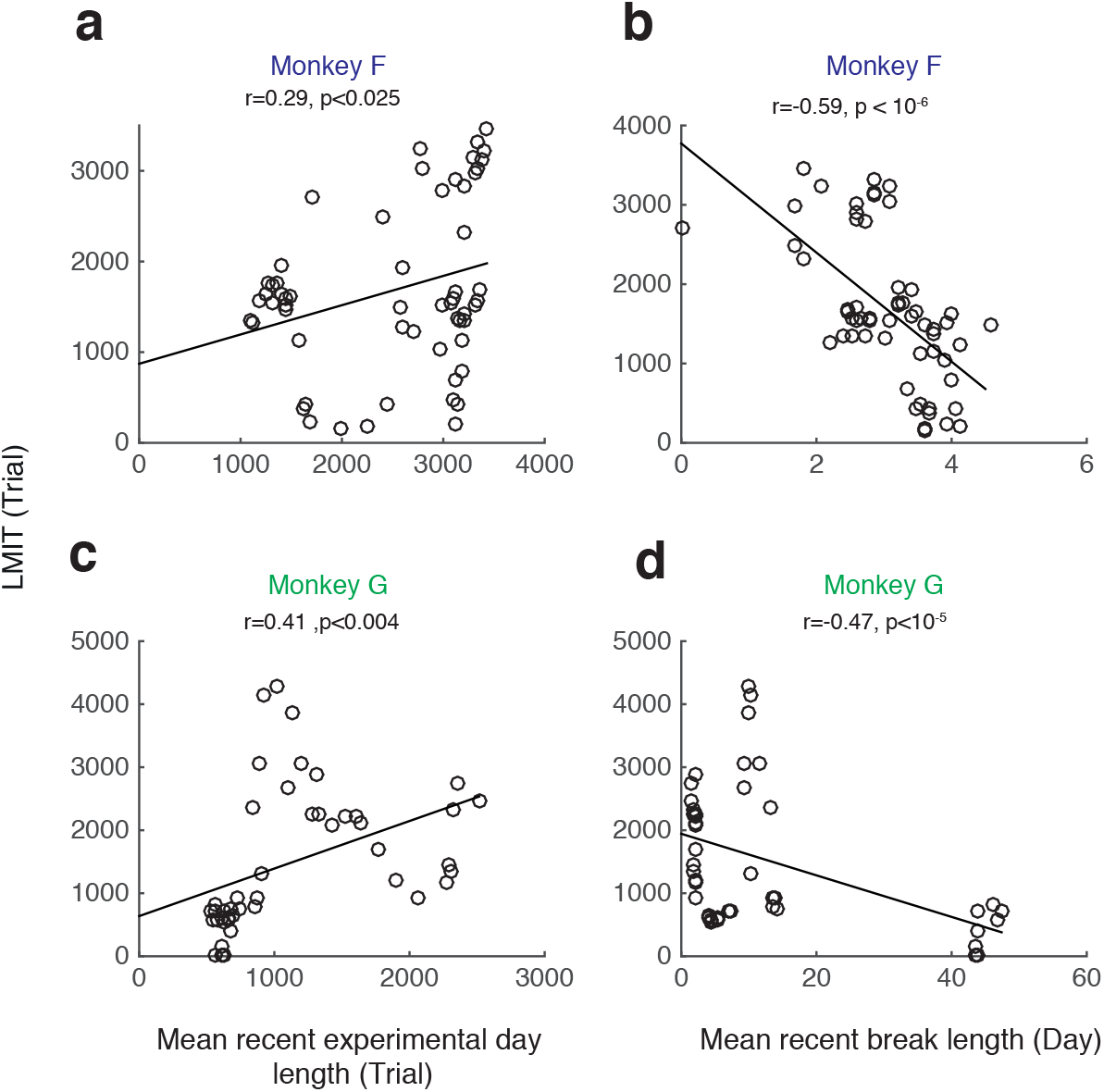
Data shows that the LMIT is affected by the *intensity* of recent consecutive experimental sessions. (**a,c**) LMIT increases as the monkeys completed more trials in recent days. The LMIT and the mean recent experimental day length (how many trials monkeys completed in one experimental day) show a significant positive correlation [permutation test: *p* < 0.025 for Monkey F and *p* < 0.004 for Monkey G]. (**b,d**) LMIT decreases as monkeys recently have had longer holidays (characterized by the mean recent break length, which is defined by the mean inter-experimental-interval). The LMIT and the mean recent break length show a significant negative correlation [permutation test: *p* < 10^−6^ for Monkey F and *p* < 10^−5^ for Monkey G]. The LMIT is computed in the same way as the previous figures with sliding windows, and both the mean recent experimental day length and the mean recent break length are the average of past 12 experimental days of each data point. Please see Figure S4 for the time course of changes in these variables, and also Figure S5 for the correlations with *w_Slow_*, instead of LMIT.

## 3 Discussion

Deviations from the matching law are sometimes interpreted as failures due to limitations of our cognitive or perceptual systems, such as neural noise. We showed that they may actually reflect a sophisticated strategy to deal with the variability and unpredictability of non-stationary environments. The decisions of the monkeys seem to be based on reward integration on multiple timescales, some of which are surprisingly long (multiple sessions). The idea of learning on multiple timescales echoes with previous observations on the behavior of primates [11, 14] and pigeons [2], as well as other computational model studies [54, 26, 21]. Our work also echoes with previous studies on suboptimal perceptual decisions [56, 4, 1]. Moreover, in the recent years there has been accumulating evidence that several neural systems – ranging from individual neurons studied in vitro, to complex neural circuits studied in vivo – have the ability to integrate their inputs, including those that represent rewards, on multiple timescales [48, 38, 27, 58, 33, 55, 10, 8, 24, 7, 42, 60]. Our results provide new insights into the computational advantage of biological processes that operate on multiple timescales.

Integrations over multiple timescales can be implemented in several ways. Although it is difficult to determine the exact mechanism, we can make some general considerations about the properties that this mechanism should have. In our model the effective time scale over which rewards are integrated is modulated by changing the relative weights of multiple integrators that operate on diverse timescales. We show in the Supplementary Material that a model with a single varying timescale would be incompatible with the data (Figure S6). This indicates that there must be processes operating on multiple timescales and that the final choice is the result of non-trivial interactions between these processes. This is not too surprising as there are significant computational advantages when synapses are endowed with metaplasticity [15, 21, 6], or when memory systems are partitioned in interacting subsystems that are responsible for preserving memories on different timescales (see e.g.[44]). Both of these cases show that multiple timescales are important for memory consolidation, and memory is certainly a fundamental component when integration over time is required. In fact, our simulation studies show that the previously proposed model of synaptic metaplasticity [15, 21] can capture some of the key aspects of our data [23] (See Figure S7).

We used our simple model to study the effects on decision making of changes in the relative weights of different integrators that operate on multiple timescales. Here we did not model explicitly the mechanism which controls and tunes these relative weights, which is a difficult and interesting problem (see [21], Supplementary Material S2 and Figure S7 for a possible implementation by a synaptic metaplasticity model). We also did not analyze transients induced by sudden changes in the statistics of the choice values (which occurs at every block change), as we focused on the effect of reward history on a timescale that is longer than the block size. We and other researchers have previously looked specifically at the mechanism which may speed up the convergence to more accurate estimates when a change in the environment is detected [39, 5, 35, 18, 34, 59, 21, 53, 40, 61]. These mechanisms are probably complementary to those that we studied, and in the future we will try to combine all these mechanisms in a unified model.

We showed that the harvesting performance of the monkey increases when the relative weights of the integrators are appropriately modified. This improvement could be significantly larger in other situations. In a two-choice task the harvesting performance varies in a rather limited range when the behavior goes from random (or when the monkey responds in any other way that completely ignores the reward ratios) to optimal. This is a well-known limitation which makes it difficult to establish how close the behavior is to optimal. In more complex, but perhaps more realistic, tasks that involve multiple choices and timescales, the situation could be drastically different. Imagine for example a task in which there are 100 different choices and only one of them is rewarded with a probability that is significantly different from zero, using a schedule similar to the one of the two-choice task that we analyzed. The rewarding targets may change time to time; but within a restricted fraction of targets, say 10 targets, whose locations also change on a slower timescale. In this case the long timescales contain important information about the possible rewarding targets that should be considered. Ignoring this information would lead to a significant decrease in the performance (see Fig. S8).

How and why animals show matching behavior has been of great interest in psychology, economics and neuroscience [43, 32, 47, 31]. While the matching law was classically studied in stationary environments [19], matching has been shown to be a good description of behavior even in non-stationary environments [52, 28] when one considers relatively short timescales. In fact, if one fits a reward integrator model characterized by a single timescale to the data that we analyzed, the estimated timescale would be approximately the optimal one [52]. Additional analysis, however, has suggested that at least two, relatively short, timescales (approximately 2 and 20 trials) are required to characterize the reward integrators. The mechanism that we studied operates on much longer timescales and probably complements those that have been analyzed in the past. The integration timescales that can be detected with the LMIT analysis are surprisingly longer than what have normally been considered (including in [52, 11]), and they vary over time on even longer timescales. Our analysis cannot determine whether the LMIT changes because the relative weights of different integrators are modified [11, 15, 21, 6], or because the intrinsic time constants of the integrators change. This is because the timescale that we consider is extremely long. It is very possible that both types of models of the LMIT can accurately describe the data and capture the correlations between the LMIT and undermatching. Moreover, it is difficult to study the interaction between the mechanisms that we studied and those that determine the optimal short integration timescales. It is possible that the long timescales make the effective timescales of the fast integrators longer, but it is also possible that the mechanisms implementing the two types of integrators are completely independent and that they both contribute to the final harvesting efficiency.

The idea of bias variance tradeoff that we discussed in this paper has been extensively studied in machine learning [13]. When fitting a model to data, the ideal model should both accurately capture the observed data, and also generalize to unseen data. One way of improving this problem is to introduce a prior belief about the data into the model, so that the model’s estimate does not entirely rely on the current observation. This idea echoes with our present findings in monkeys, as learning over a long timescale can naturally be interpreted as building a prior (see Supplementary Material S1 for more mathematical discussions).

Our results also provide a new insight into well-documented, suboptimal, exploratory choice behaviors in animals, in relation to what is often refereed to as the exploitation-exploration tradeoff [9, 12]. It has been reported that animals often fail to exploit the optimal choice but instead show matching-like, more random, behavior in simple armed bandit tasks (e.g. [41]). Computationally, this has been often accounted for by adding noise to the value computations or decisions. Our current study, however, suggests that such apparent sub-optimal exploration behaviors can be driven by a very slow learning, or reward memory on a very long timescale. Thus our model predicts that such exploratory behavior can be manipulated by changing the reward statistics on a long timescale, and possibly by directly stimulating the neural circuits that are responsible for the reward learning on a long timescale [25, 24, 10].

## Acknowledgement

We thank P. Dayan, L.F. Abbott, K.D. Miller, C. R. Gallistel, and K. Lloyd for fruitful discussions. Supported by the Gatsby Charitable Foundation, the Simons Foundation, the Schwartz foundation, the Kavli foundation, the Grossman Foundation, the Israel Science Foundation (Grant No. 757/16), the National Eye Institute and the Howard Hughes Medical Institute.

## Methods

### Subjects and the task

As the full details of the experimental protocols are reported in [52, 11], here we only provide a brief description. We used two adult male rhesus monkeys (Macaca mulatta) weighing 7 and 12 kg. To initiate a series of trials, the animals had to fixate a black central fixation cross for a period of 300 ms. This was followed by a presentation screen, consisting of the black central fixation cross and two choice targets at a pair of mirror symmetric locations in opposite hemifields – a green circle and a red circle. The animals had to fixate a central point during this period (1–2 s). The location of the red target and the green target was counter balanced across trials. At the end of this delay period, the fixation cross became white (Go cue), signaling that the animal should indicate its choice with an eye movement to one of the two choice targets within a 1 s grace period. The animal was required to maintain its gaze on the chosen target for a further variable hold period of 300-600 ms. If the chosen target was ‘baited’ at the time of the animal’s choice, a fixed magnitude fruit juice reward was delivered during this hold period. At the end of the hold period the presentation screen reappeared, cueing the animal to return its gaze to the fixation cross within a grace period of 1 s in order to trigger the onset of the next trial. Trials continued as long as the animal maintained its gaze within a 2-degree spatial window centered on the location of the fixation cross or chosen target. A failure of maintaining a gaze lead to a 2–4 sec of a timeout period, before the animal was once again given the opportunity to fixate the central cross.

The sum of the reward-baiting probabilities on the two colored targets was set constant at approximately 0.35 rewards per trial. An empty target was baited with the given baiting probability. When a target was baited, a reward was delivered if the animal chose the target. This made the target empty, with no reward available for the target, until the target became baited. The relative reward-baiting probabilities on the two colors were set constant for a block of trials and changed without a signal between blocks of trials. The typical block size was about 100 trials. On each block the relative baiting probabilities on the two colors was chosen unpredictably from a subset of the ratios (though not all experiments used all ratios). Each session consisted of multiple blocks of trials, and animals performed typically several sessions per an experimental day. Experimental days were separated in a non-systematic way.

Throughout the experiments, a “changeover delay” (COD) was imposed. This is a common manipulation necessary to ensure matching behavior by discouraging simple alternating strategies

[52]. We implemented a COD by delaying delivery of programmed rewards following switches between colors until the second consecutive choice of the new color. Monkeys learned the COD: they chose the new color twice in a raw with a probability of over 95%. As second choices after switches were imposed by the COD, we excluded those choice from our analysis. We called the remaining choice as ‘free choice’. In practice, this was achieved by treating the first two choices that mark a switch between colors as a single choice.

### Defining and measuring undermatching

There are two different choice biases that we used in our analysis: the undermatching and the color bias.

Undermatching, or deviation from matching behavior, was defined by 1 minus the slope of block-wise matching behavior. The slope of block-wise matching behavior was estimated by regressing a block-by-block fraction of choice allocated to the green target to the fraction of rewards collected from the target in a corresponding block.

While the undermatching was defined as a bias in slope, the color bias was defined by the intercept of matching behavior. Specifically, we defined the choice bias in color as the value of the fitted line at a reward fraction of 0.5. We used this bias when estimating a long timescale of reward integration (see below). For simplicity, we call ‘1-slope’ as bias or under-matching, while intercept as intercept or color bias.

### Description of the model and its simulations

We extended a previously introduced model [52] in which the income for each target (*I_G_* and *I_R_*, for the green and red target respectively) was integrated on a single timescale. In our model we consider multiple timescales as follows. The probability of choosing the red target is:

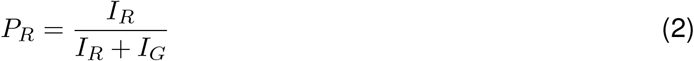

In [52], the local incomes are assumed to be computed on a single timescale. Here we assume that the incomes are computed on multiple timescales in parallel:

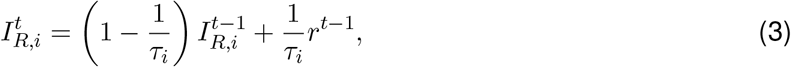

where 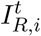 is the local income from target R on trial *t* (*t* = 1, 2, 3, …) computed over the timescale of *τ_i_*, and *r*^*t*−1^ is 1 (0) when the target was rewarded (no-rewarded) at *t* − 1. The local income is a weighted sum of different timescales:

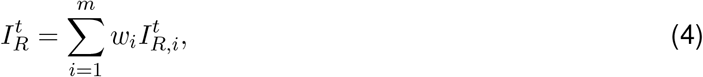

where the weights *w_i_*’s are normalized so that *w*_1_ + *w*_2_ +.. = 1. In the model simulated in Figure 2, the parameters were *m* = 2, *τ_1_* = *τ_Fast_* = 5 trials, *τ_2_* = *τ_S1ow_* = 10, 000 trials.

We computed the fluctuation of Monkey’s choice in Figure 3d as follows. First, monkey’s choice time series was smoothed via two half-gaussian kernels with standard deviations of *σ* = 8 trials and *σ* = 50 trials, with a span of 200 trials. This gave us two time-series, a fast one with *σ* = 8 and a slow one with*σ* = 50, which we used to compute the bias and variance. We defined the fluctuation as the standard deviation of the fast one over the slow one. The bias was defined by 1– the slope of block-wise matching behaviors. In Figure 3de, the model with different *w_S1ow_* was simulated in Monkey F’s reward schedule. *τ_Fast_* = 2, *τ_S1ow_* = 1000 trials.

### The longest measurable integration timescale (LMIT)

The aim is to determine whether past biases in rewards could affect the present choice of the animal. We computed the lagged cross correlation between reward bias and choice bias. The reward bias over *N* trials is defined as 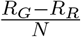, where *R_G_* (*R_R_*) is the total number of rewards that were collected from Green (Red) and *N* is the total number of trials. The choice bias is the intercept *y* with *x* = 0.5 of the linear fit of matching behavior. Then we took a sliding reference window of 25 sessions. For each sliding window, we computed the correlation between choice bias computed over 25 sessions and the reward bias computed over a 25 sessions lagged in time (lag = 0, −1, −2, −3, −4, −5 sessions). These correlations were fitted by weighted least squares 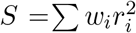 with weights *w_i_*, = γ^*i*^ where *i* = 0, 1, 2.. was the lag, as the correlations were more reliable for smaller lags. We used γ = 0.5 but we found that our results were robust against changes in γ. We defined the point in which the fitted line crosses the time lag axis (or 0 correlation) as the raw maximum correlation lag. Then the LMIT was then expressed as this lag multiplied by the mean session length (in trials) of 25 sessions in the reference window. The significance of the correlation between under-matching and the LMIT was determined by a conservative piecewise permutation test which supposedly destroys the original correlations. More specifically, we considered blocks of 5 consecutive sessions and shuffled the order of blocks without perturbing the order of sessions within each block. This allowed us to create shuffled data with the original long timescale correlations. This was because observed LMITs were normally within 5 sessions. Figure 5c is smoothed by computing a moving average of 10 windows but figure 5d is raw data. First 25 sessions were excluded in order to discard potential effects due to training.

## Supplementary Information

### S1 Probability estimation of binary sequence over blocks of trials

Here we illustrate our analytical results of the probability estimation by a model that learns inputs over multiple timescales. For this, consider the following statistical estimation task. An observer (a monkey statistician) observes the time series, s_t_, of the (conditionally) independent drops of a coin (heads *s* = 1, tails *s* = 0), where the head probability, *p*, is constant over each block of *T* coin drops (trials), while the probability changes on the first trial of each block to a new *p*, by independently sampled from a distribution, *π*(·), on the interval [0, 1] (For now, our default choice will be the uniform distirbution on [0, 1]). The observer’s goal is to estimate the current head probability, *p*, by collecting/integrating the observations over trials, over a flexible combination of two, fixed, time scales. More precisely the current estimate, *v_t_*, of *p*, at time *t*, is given by

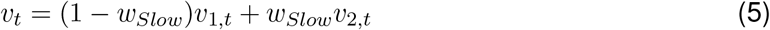

where 0 ≤ *w_S1ow_* ≤ 1, and *v*_1,*t*_ and *v*_2,*t*_ are leaky integrations of the recent history of *s_t_* over the time scales *τ_1_* (fast) and *τ_2_* (slow), respectively, i.e.

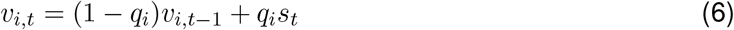

where the learning rates, *q_i_*, are defined by the inverse of time constants 1/*τ_i_*. Solving (in steady state, i.e. after many blocks) we have

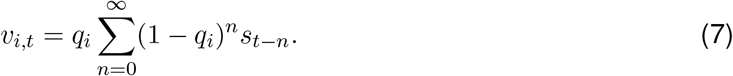

We assume that the time-scales, *q*_1_ and *q*_2_, are fixed, perhaps reflecting a form of hardware constraints, but we consider *w_S1ow_* to be flexible. We want to find the optimal *w_S1ow_* leading to the minimum possible average square error for the estimator *v_t_*, i.e. the *w_S1ow_* that minimizes the long-time average of (*v_t_* − *p*)^2^, given the knowledge of block size *T* and the internal time-scales, *τ_i_*.

We will adopt the following index notation. We use *t* as the trial index, *n* as the trial lag (into the past), and *k* as the block index. We denote the current block by *k* = 0, with *k* = 1, 2, … indicating past blocks (so k > 0 indicates the block-lag into the past). Thus *p*_0_ is the head probability of the current block, *p*_1_ that of the previous block, and so on. We choose the time origin such that the first trial of the current block (*k* = 0) has *t* = 1, with trials in past blocks having zero or negative *t*’s.

We also adopt the following averaging notations. We denote the average of a quantity, conditional on knowing the full sequence of block-probabilities *p*_0:∞_, by 〈·〉, i.e.

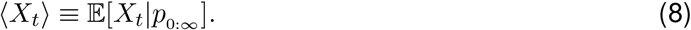

We indicate averaging over *p*_0:∞_ by [·]_π_, i.e.

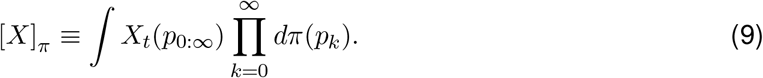

Finally we indicate averaging over the duration of a block by a bar:

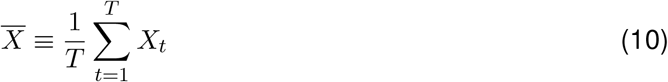

Thus we set out to calculate the long-run average square error, which in the above notation is given by

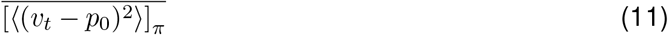

and then find the optimal *w_S1ow_* that minimizes this cost.

We start by evaluating ((*v_t_* − *p*_0_)^2^) which can be decomposed in the standard way, into the variance of the estimator and its squared bias, i.e.

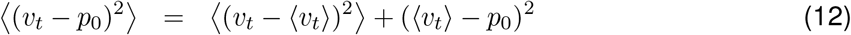

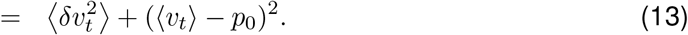

In model-fitting and parameter estimation, variance generally quantifies the degree to which an estimator is sensitive to noise in the data (the noise in {*s_t_*}, in our case), in other words it quantifies how much it can fit noise, i.e. over-fit, while the bias normally quantifies how rigid or inflexible a model is (its inability to conform to and capture certain aspects of the data, hence creating biases). Generally speaking, the more complex a model, the more flexible it is, and the lower is its bias and higher its variance (it is prone to over-fitting). One way of reducing the variance and estimator-flexibility is to introduce a prior (which could represent past experience), making the estimator not rely entirely on currently observed data but also on prior knowledge (or past experience). In our case, the integrator with the slow time-scale, *v*_2,*t*_ serves this purpose; it represents the prior knowledge acquired over long time scales (previous blocks), and is less altered by observations in the current block (it has a smaller learning rate). But by that virtue it is less flexible and retains its bias longer and is slow in adapting to the present value of *p* in the current block.

To summarize

- bias = prejudice = rigidity = low model complexity = slow time scale,
- variance = open-mindedness = flexibility = high model complexity = fast time scale.

A good model/observer must balance the two.

#### S1.1 Variance

We will first look at the long-run average variance 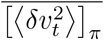. From Eq. (5), the conditional variance is given by

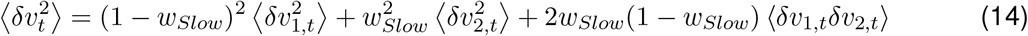

where *δv_i,t_* ≡ *v_i,t_* − 〈*v_i,t_*〉. From Eq. (7), *v_i,t_* is a linear combination of independent random variables, *s_t_* (the latter are independent only when conditioning/fixing *p*_0:∞_), thus its variance is the sum of the variances of the terms in this sum. The variance of *s_t−n_* in block *k* is given by

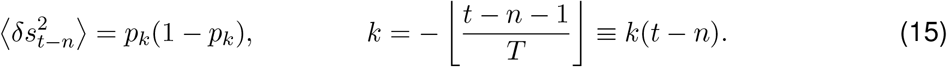

Thus

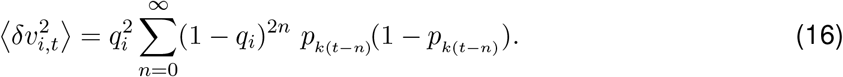

Similarly

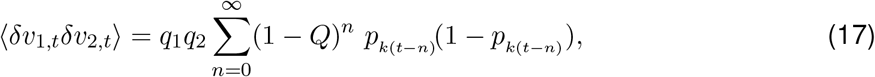

where we defined

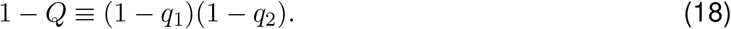

Since [*p_k_*(1 − *p_k_*)]_π_ is the same in all blocks, hence independent of *k*(*t*−*n*), we can readily calculate the variance averaged over *p*_0:∞_, by summing the infinite geometric series, obtaining

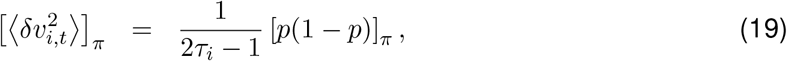

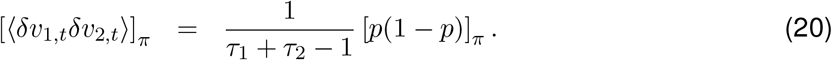

where we used 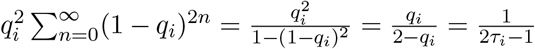 and 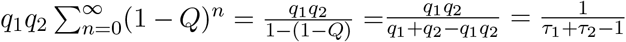. In particular, 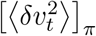 is time independent:

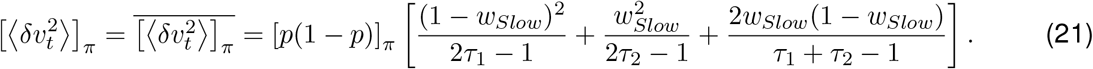

##### S1.1.1 Variance conditional on *p*_0_: transient behavior

For completeness, we will also calculate the variance conditional on *p*_0_ as well, obtaining its full transient behavior throughout the block. That is, here we will only average over *p*_1:∞_, but not over *t* and *p*_0_. Going back to Eq. (16), we rewrite it by decomposing the sum into sums over blocks:

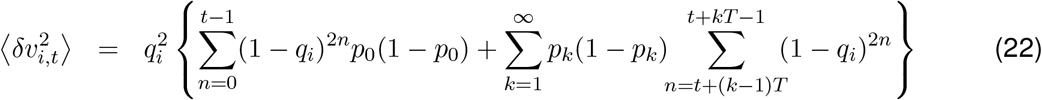

It helps to rewrite this in the form

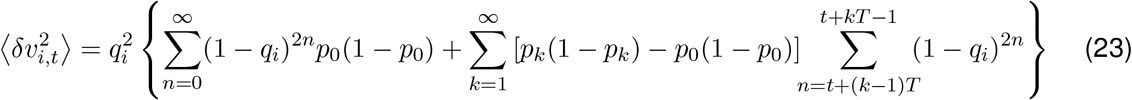

where the first term pretends that the probability was *p*_0_ in the entire past, and the second term corrects for this by adding the difference of the variances accumulated over previous blocks contributed by the true probability, *p_k_*, and the current one, *p*_0_, respectively. By the geometric series formula the sum over block *k* is given by

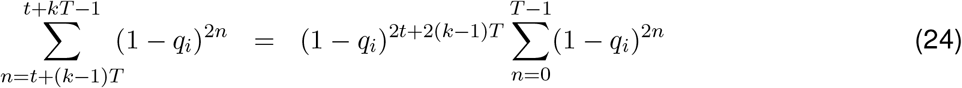

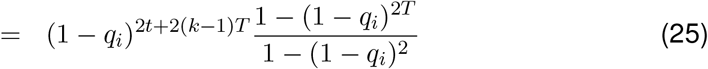

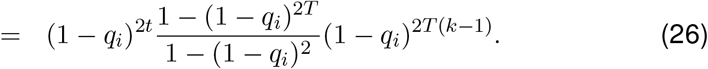

Using 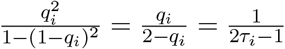, we then have

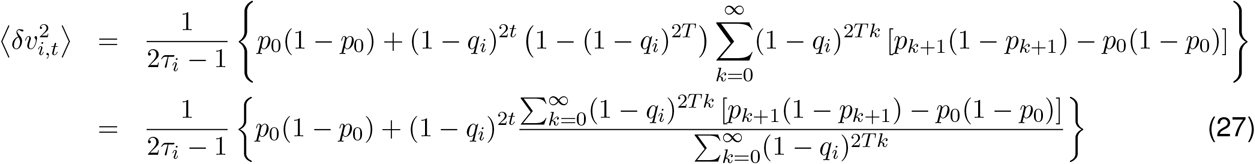

Similarly for (*δv*_1,*t*_*δv*_2,*t*_) we have

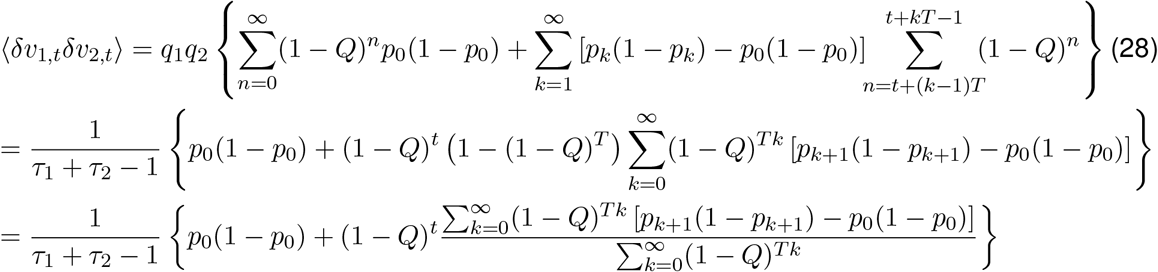

where we used 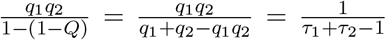. Averaging over *p*_1:∞_ and summing the infinite geometric series over blocks, combining contributions as in Eq. (14), and using Eq. (21), we then obtain

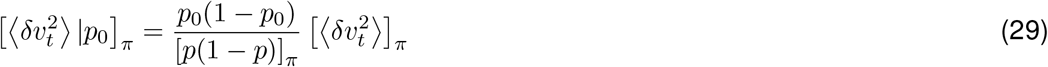

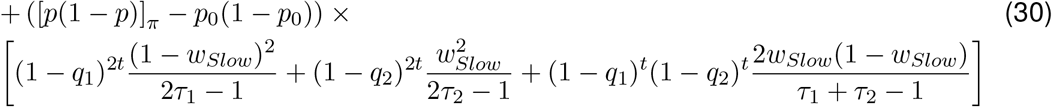

Here, the first line gives the steady state value of the variance in the current block if it was infinitely long, and the second and the third line gives the transient memory of variance from previous trials, which wears off for *t* ≫ *τ*_2_. It starts from a value equal to the average variance 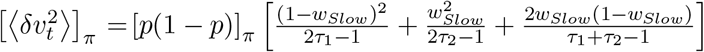 at *t* = 0 and eventually (given an infinitely long current block) relaxes to its steady-state value based on the current *p*_0_ as opposed to the average, i.e. to 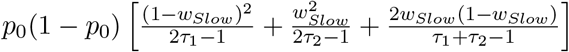.

#### S1.2 Squared bias

First let us calculate 〈*v_i,t_*〉. Using the notation of Eq. (15), from Eq. (7) we have

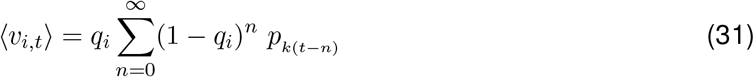

Again we can decompose this over blocks:

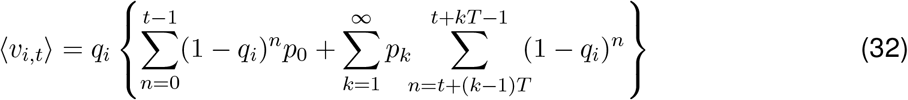

and again it helps to rewrite this as

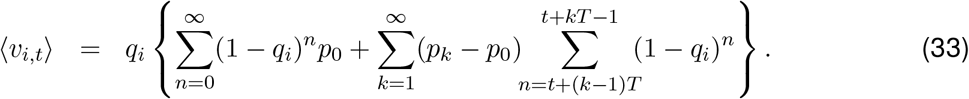

Noting that 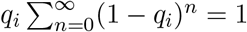, for the bias component, *b_i,t_* ≡ 〈*v_i,t_*〉 − *p*_0_, we obtain

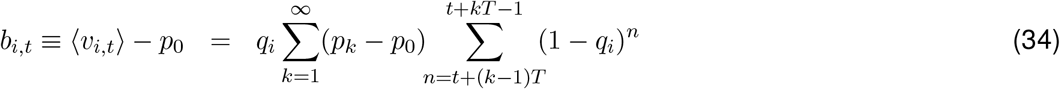

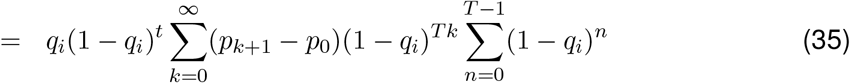

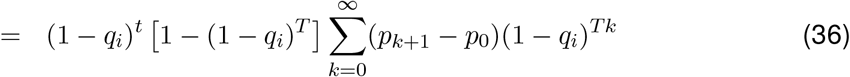

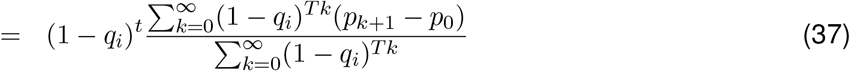

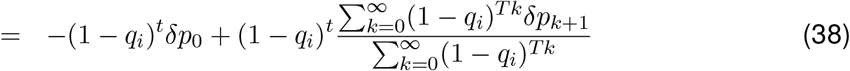

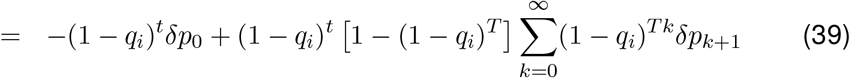

where we defined

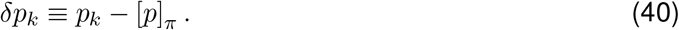

Note that we can write the bias, Eq. (39), in the form

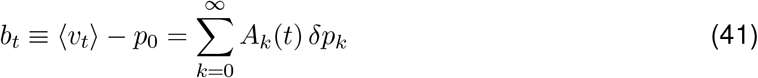

where we defined

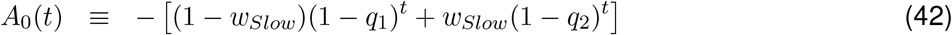

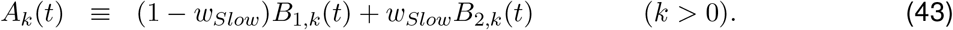

and

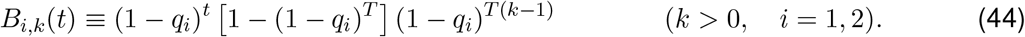

The bias squared is then given by

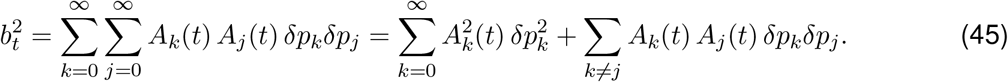

Since *δp_k_* are zero-mean independent variables, averaging over them kills the second, off-diagonal term (this is true even if we don’t average over *p*_0_) in the above expression. The bias squared averaged over *p_k_* in the previous blocks (but not on *p*_0_) is thus given by

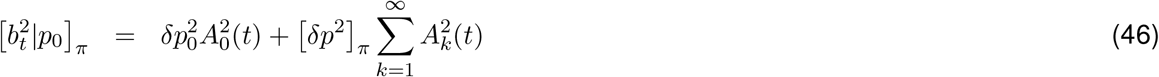

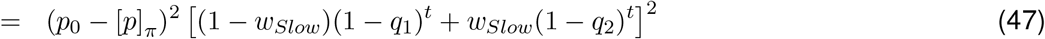

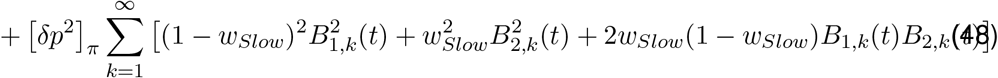

Now we have

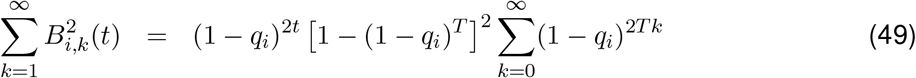

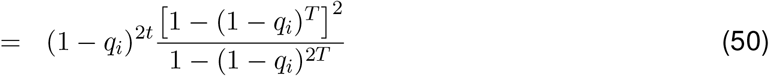

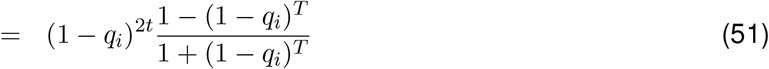

and (using 1 − *Q* ≡ (1 − *q*_1_)(1 − *q*_2_))

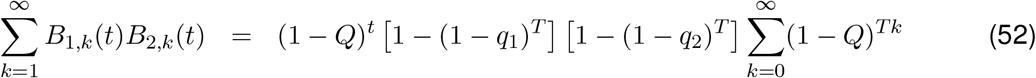

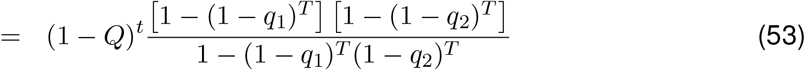

yielding

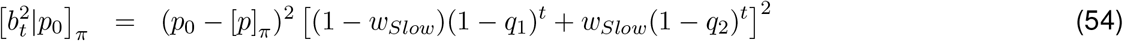

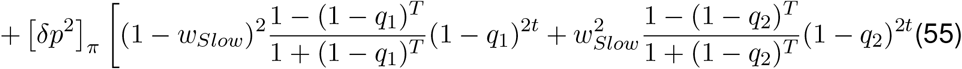

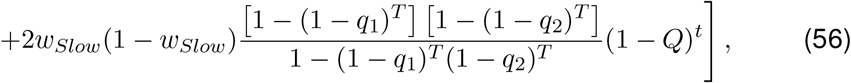

for the transient behavior of conditional average bias squared in the current block.

Averaging over p0 yields

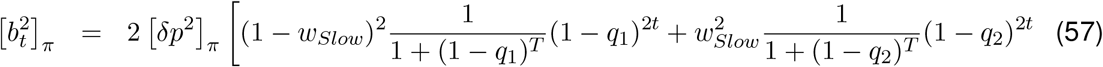

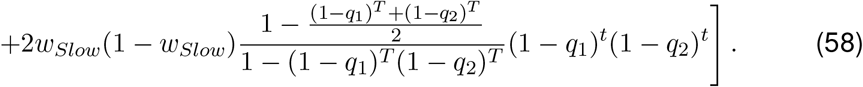

Finally, to average over *t* ranging over the block, we use

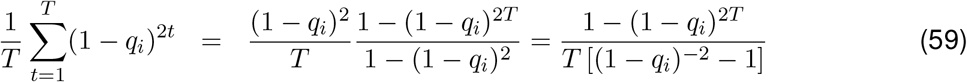

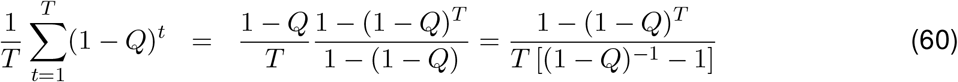

to obtain

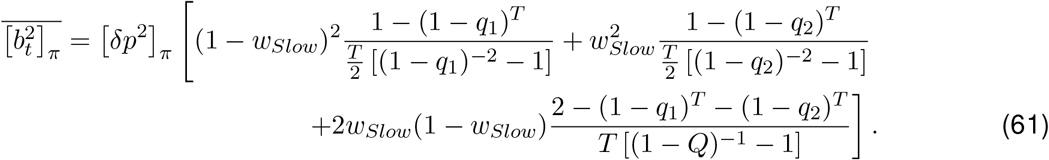

In the regime where *q*_2_ ≪ *T*^−1^ ≪ *q*_1_ ≪ 1 (or *τ*_2_ ≫ *T* ≫ *τ_i_* ≫ 1), we have approximately (1 − *q*_1_)^*T*^ ≈ 0, (1 − *q*_2_)^*T*^ ≈ (1 − *q*_2_)^*T*^ and [(1 − *q_i_*)^−2^ − 1] ≈ 2*q_i_* and [(1 − *Q*)^−1^ − 1] ≈ *q*_1_ + *q*_2_, yielding

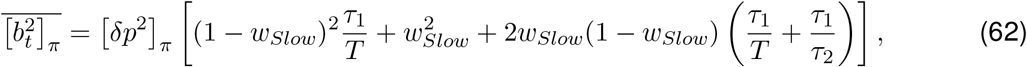

and in the limit *τ*_2_, *T* → ∞:

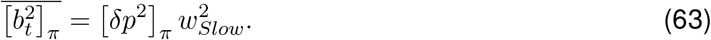

#### S1.3 Average squared error, optimal *w_Slow_*, and undermatching

The long-run average squared error is the sum of average variance and average bias squared and thus from Eqs. (21) and (61) is given by

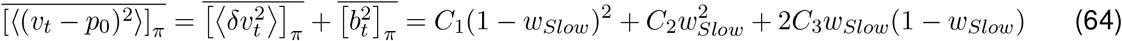

where we defined

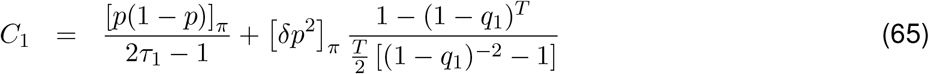

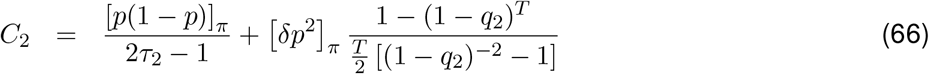

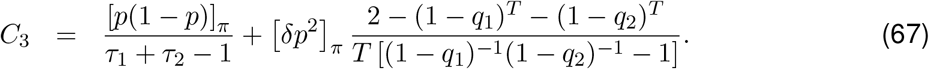

**Figure S1:**
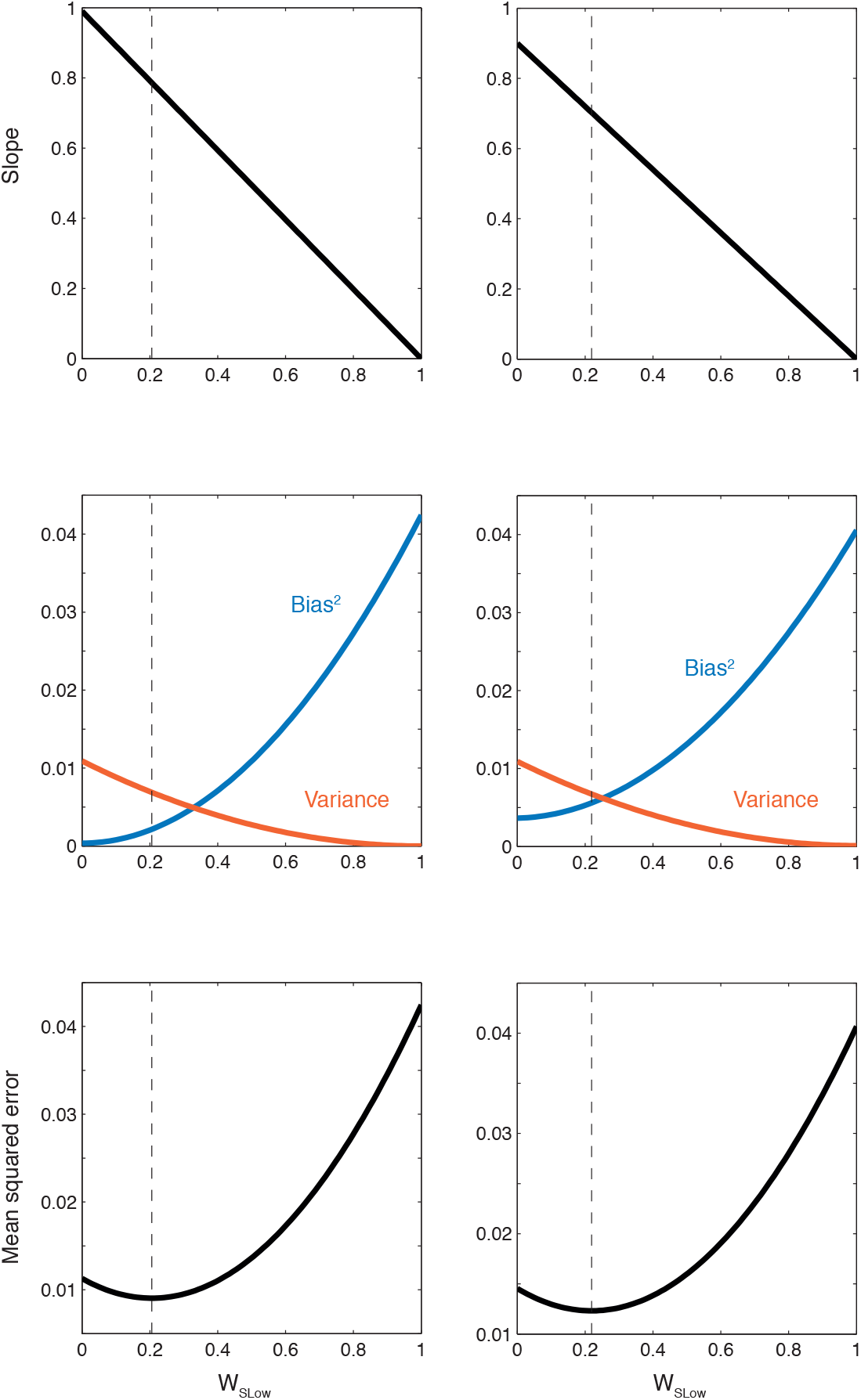
Plots of analytical results: block-averaged undermatching slope, Eq. (88) (top panels), variance Eq. (21) and average squared bias Eq. (61) (middle panels) and average squared error Eq. (64) (bottom panels). For the plots on the left, we used *τ*_1_ = 10, *τ*_2_ = 100000, and *T* = 1000, and for those on the right we used *τ*_1_ = 10, *τ*_2_ = 1000 and *T* = 100. The dashed lines show the optimal weight *w_Slow*_*, Eq. (72), in each case. Here we took the distribution π(*p*) to be the uniform distribution on [0.1, 0.9], yielding 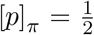 and 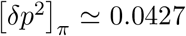 (choosing a uniform distribution on [0, 1] instead, would have yielded 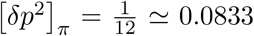, thus putting more emphasis on squared bias and shifting the minimum of bottom plots, i.e. the optimal *w_Slow_*, further to the left).

Here, in each line the first term is the contribution of the variance and the second is the contribution of the average squared bias. Note that in general

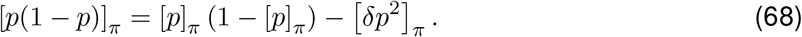

In particular, for the case where π(·) is the uniform distribution on [0, 1], we have

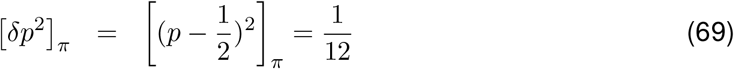

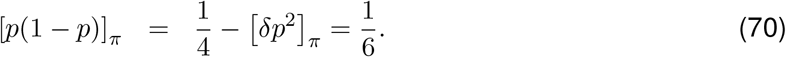

To find the optimal *w_Slow_* we have to set the derivative of Eq. (64) w.r.t. *w_Slow_* to zero. The latter is proportional to

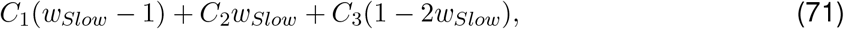

and setting it equal to zero yields

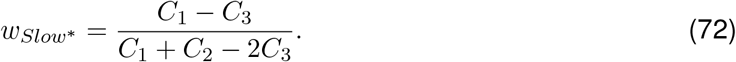

We can use Eq. (62), to simplify Eq. (65) in the regime *q*_2_ ≪ *T*^−1^ ≪ *q*_1_ ≪ 1 (or *τ*_2_ ≫ *T* ≫ *τ*_1_ ≫ 1), obtaining^1^

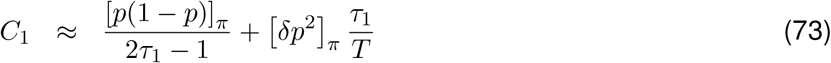

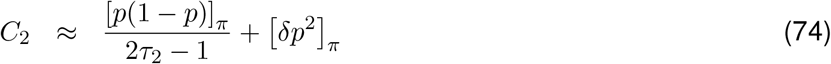

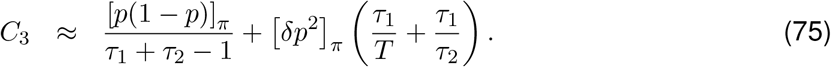

We see that the largest contribution to average error, which is *O*(1), comes from the bias squared contributed by the slow time scale (the second term in *C*_2_). After that we have the contribution of the fast time scale to variance (first term in *C*_1_) which is *O*(*τ*_1_^−1^) and smaller. For this reason, for realistic underlying time-scales, the optimal *w_Slow_*’s will turn out to mainly optimize the squared bias, and hence will be small.

It is much easier to derive these results in the extreme limit *τ*_2_, *T* → ∞ (keeping *τ*_2_ ≫ *T*). Firstly, in this case, given that *v*_2_ is a very long-term average of *s_t_*, its value is always very close to the long term average of *p*, i.e. [*p*]_π_, with small fluctuations, *δv*_2_, of the order of 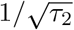. Thus we can ignore the latter and safely write

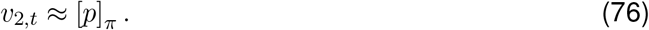

In particular, it is only *v*_1,*t*_ which contributes to the variance:

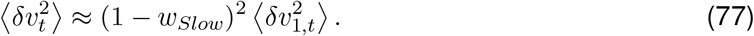

Furthermore, given that *τ*_1_ ≪ T, the main contribution to the averages of 〈*v*_1,*t*_) or 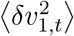 over *t* running from 1: *T* comes from *t*’s within the current block that are much larger than *τ*_1_ (i.e., we can ignore the transient behavior of *v*_1,*t*_ at the beginning of the block and only consider its steady-state behavior). This means that in Eqs. (16) and (31), we can safely replace *p*_*k*_(*t*−*n*) with *p*_0_, the head probability in the current block. The geometric series thus become infinite and we obtain

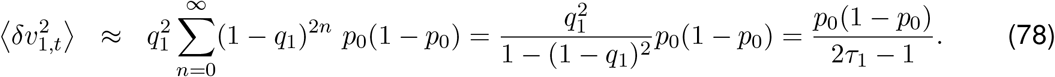

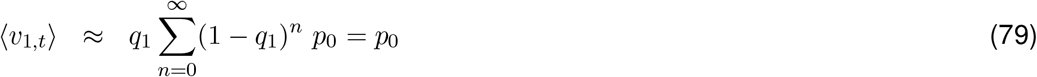

Averaging Eq. (78) over π, and using Eq. (77) we obtain

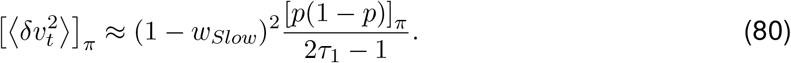

For the full bias we have *b_t_* = 〈*v_t_*〉 − *p*_0_ = (1 − *w_Slow_*) 〈*v*_1,*t*_〉 + *w_Slow_* 〈*v*_2,*t*_〉 − *p*_0_, which by Eq. (76) and (79), yields *b_t_* = *w_Slow_*([*p*]_π_ − *p*_0_) = −*w_Slow_δp*_0_ (this yields (1 − *w_Slow_*) for the undermatching slope, as an approximation to Eq. (88)). Thus

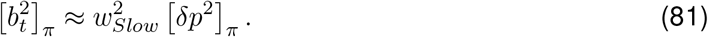

Finally for the average square error we obtain Eq. (64) with

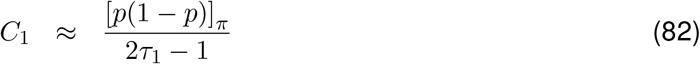

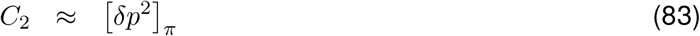

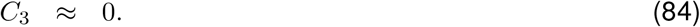

##### S1.3.1 Time-dependent undermatching slope

Going back to Eq. (39) for the bias, since the second term in Eq. (39) vanishes after averaging over *p*_*k*+1_, for the transient of bias conditional on *p*_0_ but averaged over *p_k_* in past blocks we obtain

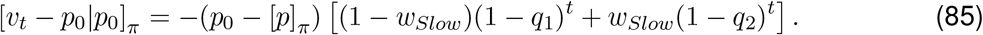

which we can also write in a form corresponding to the slop of the matching law plot

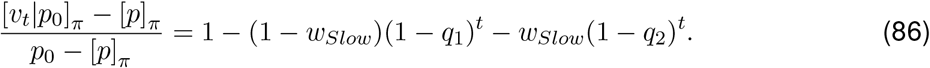

(assuming a “symmetric” distribution π(·), 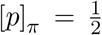. In particular, when *τ*_2_ ≫ *T* (or *q*_2_*T* ≪ 1) (1 − *q*_2_)^*t*^ remains approximately equal to unity even for *t* = *T* (at the end of the block). Thus we have

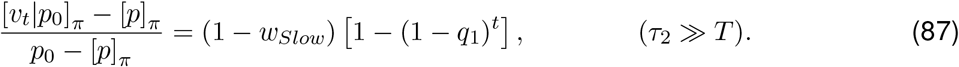

This shows that there is more undermatching at the beginning of the block, than at the end (where (1 − *q*_1_)^*t*^ ≪ 1, if *τ*_1_ ≪ *T*). If we average this over the whole block we obtain for the block-averaged (under)matching slope:

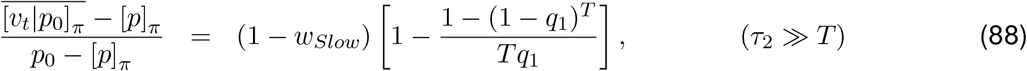

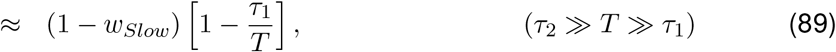

## S2 Exploring the possible neural implementation of adaptive computation: metaplastic synapses model

In our analysis, we found that monkeys integrated the reward histories over a wide range of timescales, and they changed the relative weights of the timescales to optimize the harvesting efficiency. What would be a possible neural implementation of this computation? Here we explore a possible candidate, following a meta-plastic synapses model [15] that has been recently applied to the context of decision-making [21].

Following [21], we consider a well-studied neural network model for decision-making (figure S7a) [57, 51, 14, 22, 21]. Essentially, this network produces bi-stable attractor dynamics (winner-take-all process), where each stable state corresponds to the choice of Green (when G population wins) or the choice of Red (when R population wins). Crucially, the competition is determined by the synaptic strength between the input population and the decision populations, which are trained trial-by-trial by a reward based stochastic Hebbian learning [49, 51, 14, 22, 21].

The synaptic efficacy is assumed to take one of the two strengths (weak or strong), as this follows the biophysical constraint of bounded strength. In addition to the changes in efficacy, following [15], here we introduce a metaplastic transition so that synapses can change the rate of plasticity itself [15, 21] (figure S7b). This cascade model incorporates a various chemical cascade processes taking place over multiple timescales. It has been shown that the model improves the memory performance of bounded synapses, and also the model reproduces a widely observed power-low memory decay in time. The details of implementation are described in [21]. As in [21], we assumed a ‘forgetting’ between experimental sessions which in our model is random transitions between weak and strong states between sessions.

We found that the network model captures some key aspects of the behavioral findings. We simulated our model under the same conditions as Monkey F. As seen in figure S7c, the model can reproduce the observed changes in matching behavior. In the early days of experiment, the model shows a good matching, as we assume that the synapses distribute equally over all states at the beginning of day 1. However, the distribution of synapses in the cascade model gradually shifts toward the deeper states (less plastic states). This introduces a bias toward the mean of the reward on a long timescale. Since the rewards are balanced on the long timescale, the bias is seen as deviation from the matching law, or undermatching. Figure S7d also shows that the model captures the bias variance trade-off in the data. The details of the model implementation can be found in [21]; however here we did not implement the surprise detection system, as our focus is to capture the behaviors on a long timescale.

**Figure S2:**
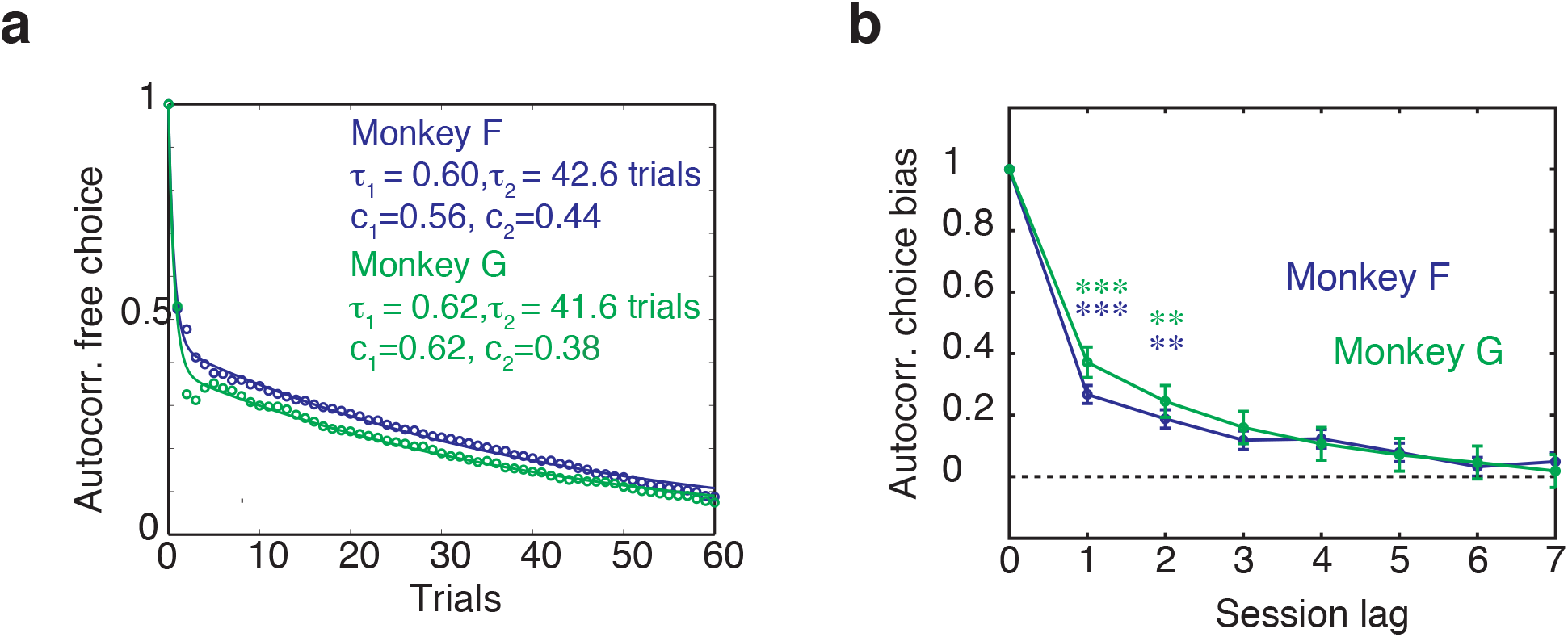
(**a**). Autocorrelation of choice history decays on multiple timescales within a block size. The solid lines indicate the fitting results with a sum of two exponents with relative weights *c*_1_, *c*_2_ and time constants *τ*_1_, *τ*_1_. Those choices reinforced by COD are excluded. (**b**) Both monkeys’ sesion-by-session choice bias show significant autocorrelations across session lags. The stars indicate the significance.

**Figure S3:**
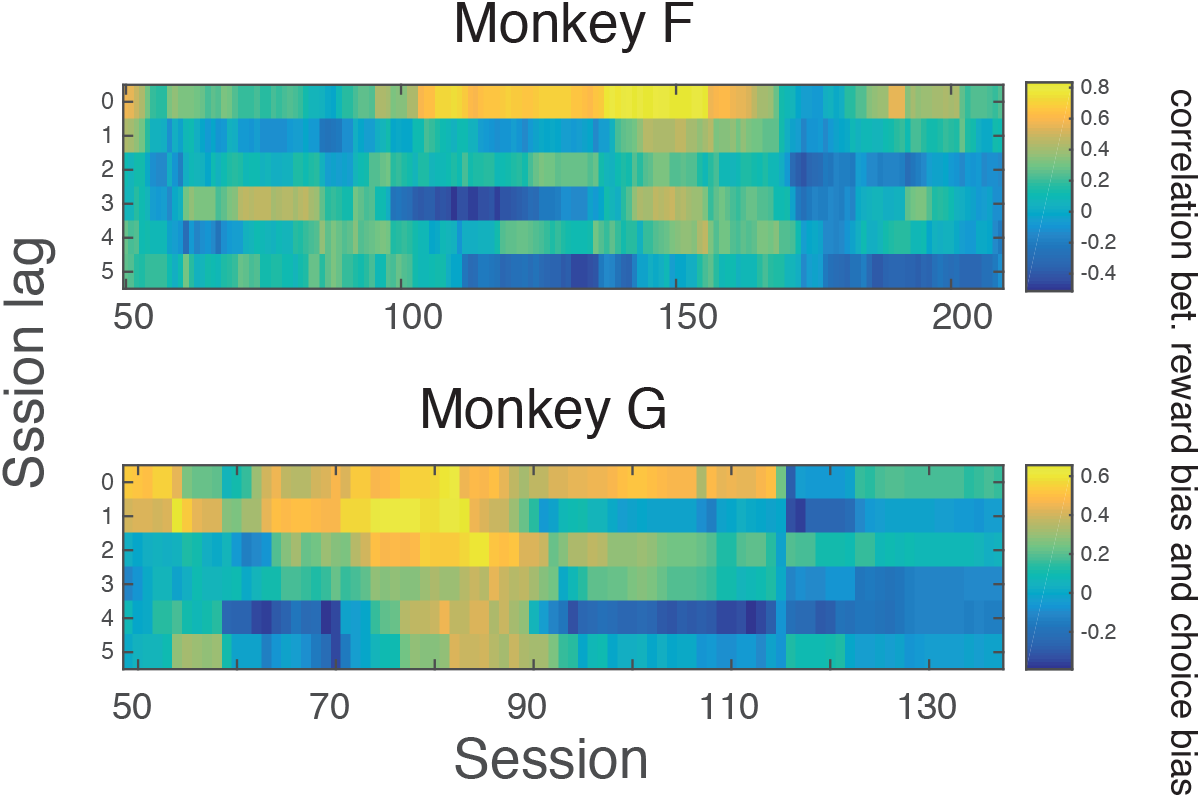
The correlation between reward and choice bias over sliding session windows.

**Figure S4:**
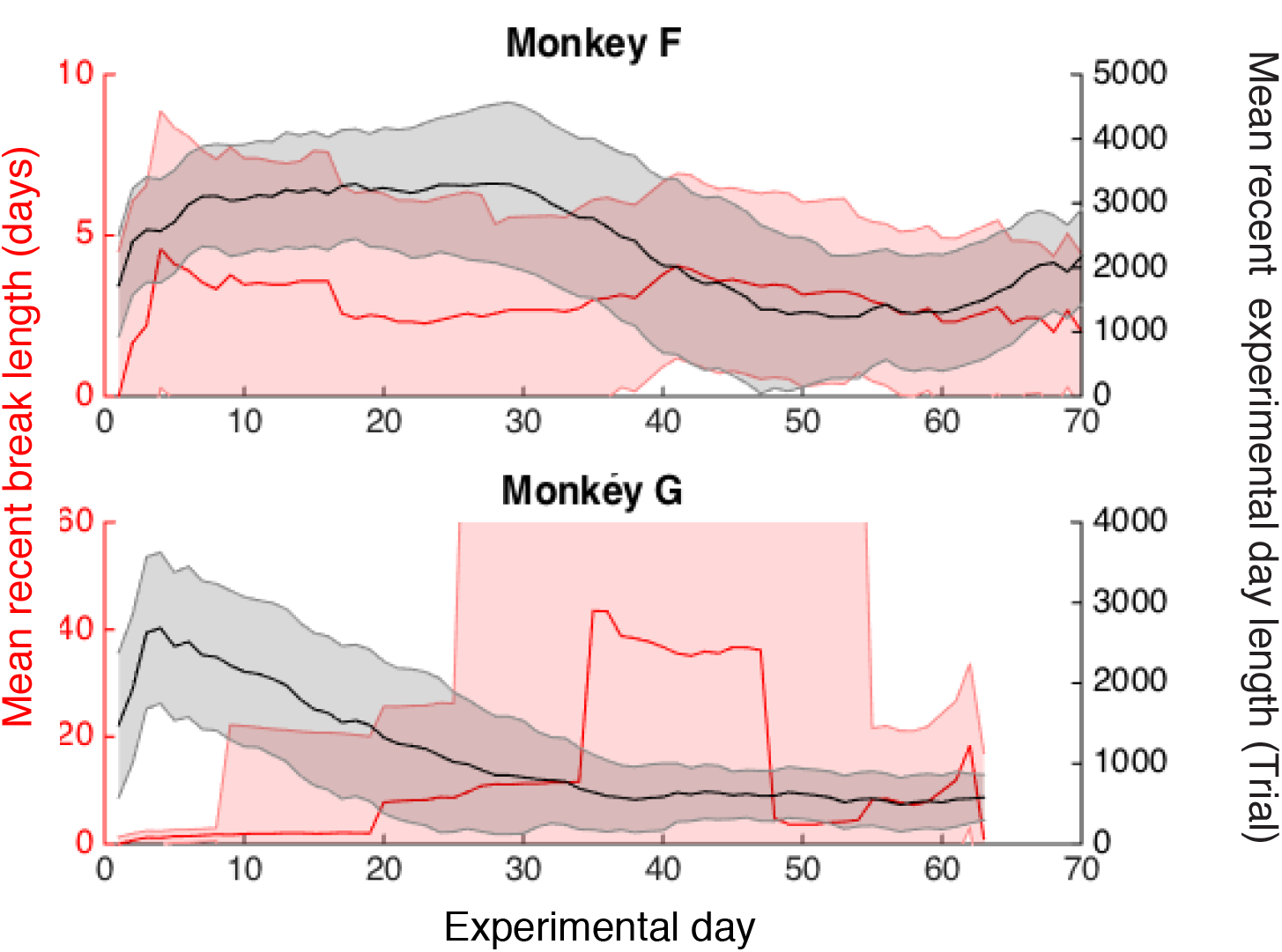
Changes in recent break length (red) and recent experimental day length (black) over the course of experiments.

**Figure S5:**
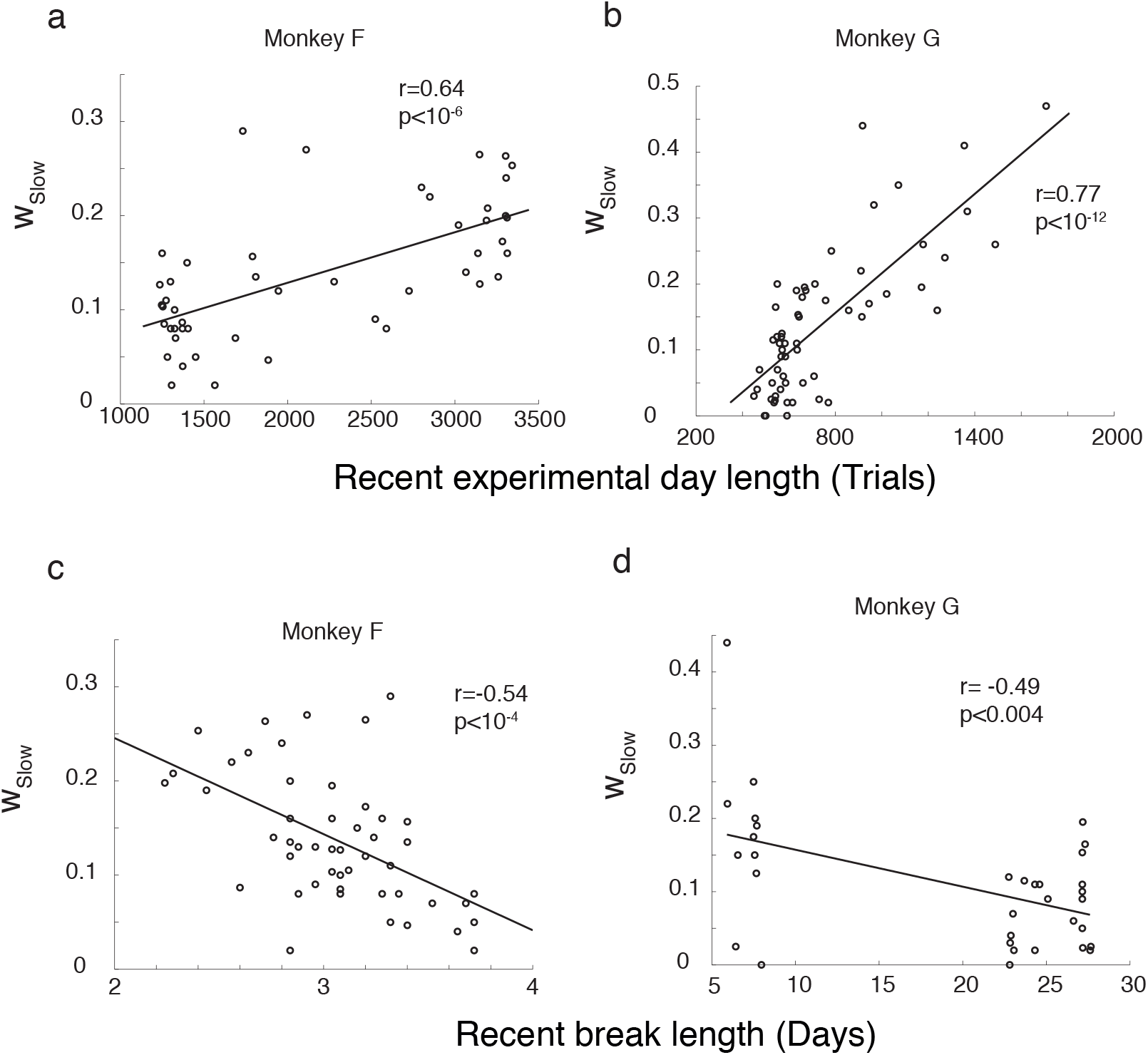
Non monotonic changes in the weight *w_Slow_* of the long timescale *τ_Slow_* reflect the experimental schedule on a long timescale. **a,b**, The weight of long timescale *w_Slow_* correlates with the recent experimental lengths. Daily estimation of *w_Slow_* is plotted against the mean length of recent experiments. The weight of the long timescale *w_Slow_* is larger when the animal constantly experienced long experiments with may trials. The mean is taken over 18 experimental days (Monkey F) and 12 experimental days (Monkey G), respectively, as they give the largest correlations. **c,d**, The weight of long timescale *w_Slow_* anti-correlates with the mean recent inter-experimental-intervals. The weight of the long timescale *w_Slow_* is smaller when the animal constantly had long inter-experimental-periods. Daily estimation of *w_Slow_* is plotted against the mean recent inter-experimental-intervals. The mean is taken over 25 experimental days (Monkey F) and 32 experimental days (Monkey G), respectively, as they give the largest magnitude of correlations.

**Figure S6:**
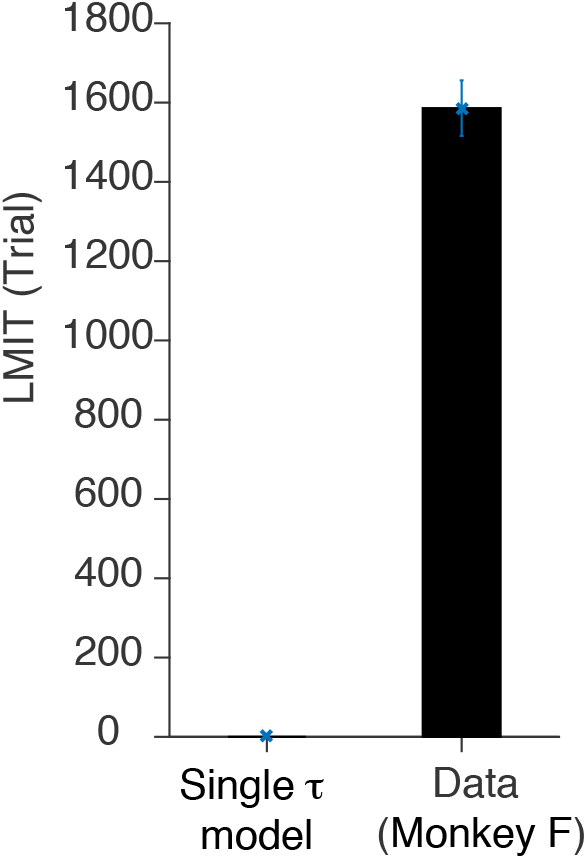
Variable-single-timescale model dose not capture data. We fitted a local matching model with a single timescale [52] to the data (Monkey F’s data). Using the fitted parameter (*τ*), we simulated the model to generate choice behaviors, which was then used to estimate the LMIT in the same manner as the other analysis. We found no LMIT from the single timescale model, because of the lack of a slow learning.

**Figure S7:**
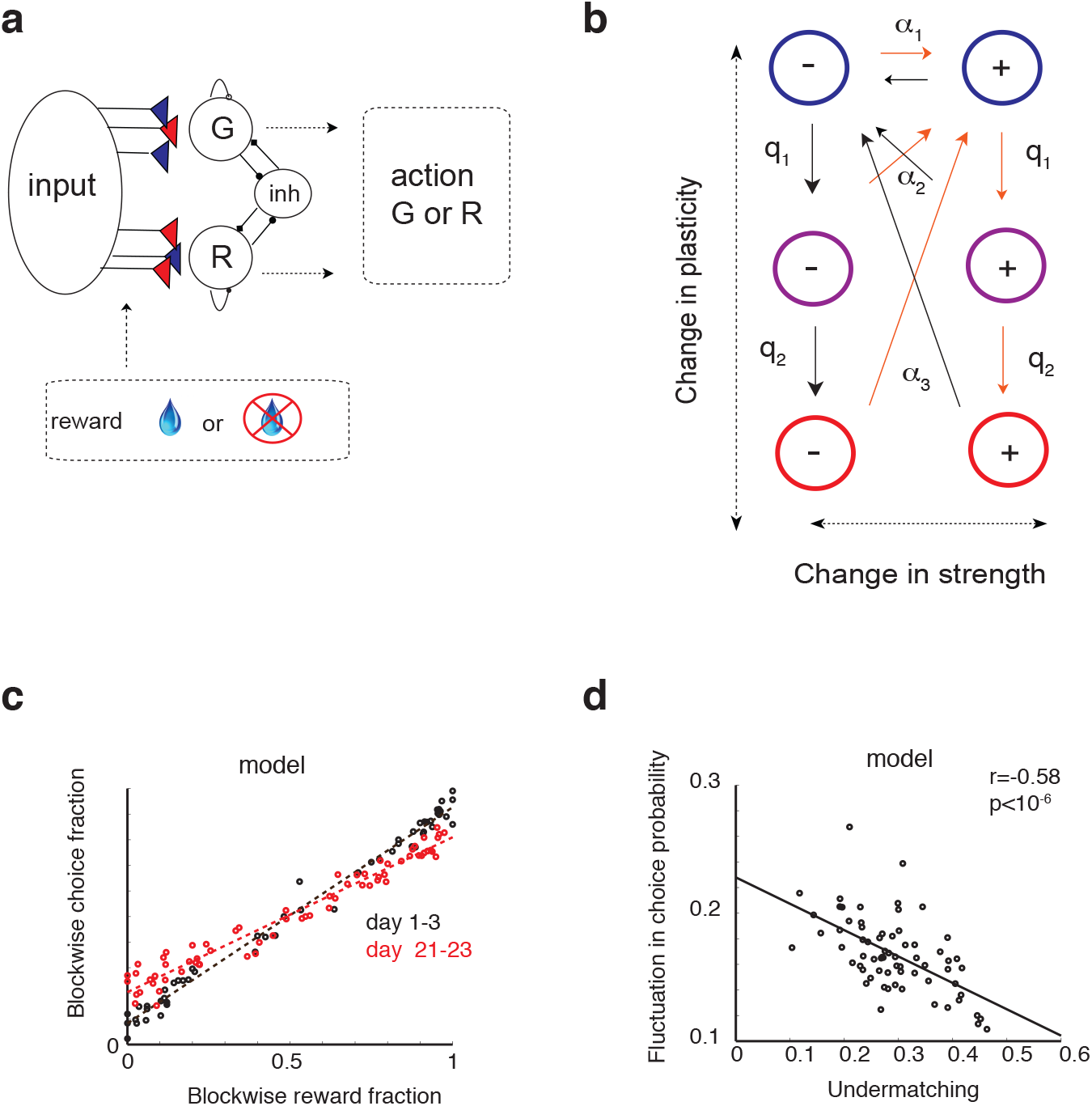
Metaplastic synapses model captures key aspects of experimental data. (**a**) Network model. The decision is made based on the competition between the two action selection populations (G,R). See [21] for details. (**b**) The cascade model of synapses. Synapses make transitions between different strength and plasticity states *α*_1_ > *α*_2_ > *α*_3_, *q*_1_ > *q*_2_. See [21] for details. (**c**) Changes in matching behavior is captured by the model. The model was simulated in the same conditions as Monkey F in data. (**d**) The model also captures the trade-off between bias (undermatching = 1–slope) and the fluctuation in choice.

**Figure S8:**
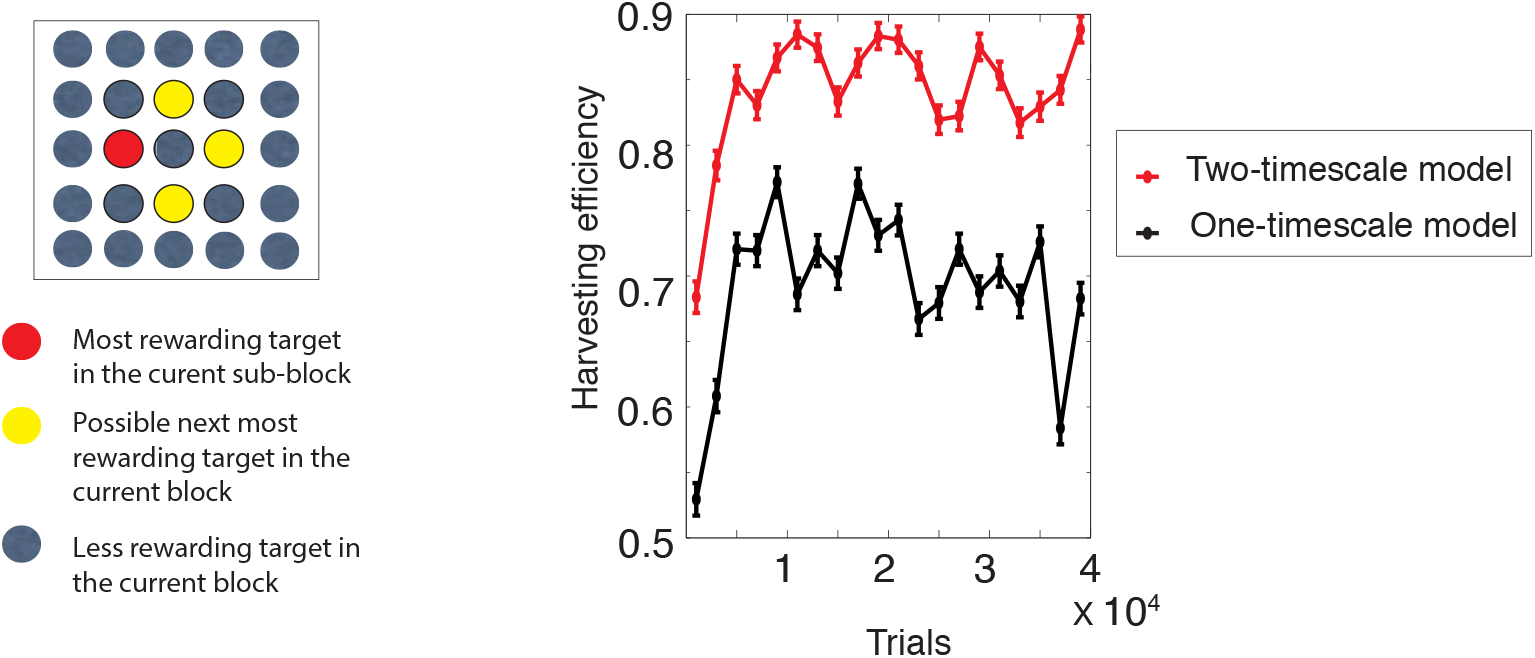
The multi-timescale model performs significantly better in a multi-armed bandit task with a hierarchical block structure. Subjects need to choose a target from 25 different options (colors are not shown to subjects). There is one target that is most rewarding in the current subblock of trials (red), and the most rewarding target would change among the ‘hot spots’ (yellow) which are fixed for the current block of trials. Crucially, the most rewarding target changes on a shorter timescale (e.g. every 20 trials), while the hot spots change on a longer timescale (e.g. every 200 trials). In such a situation, learning over a short and a long timescale is beneficial.

## Footnotes

1 To be really consistent in the apprximations, the first terms on the rights sides of Eq. (73) must also be expanded.

